# Harp: Platform Independent Deconvolution Tool

**DOI:** 10.1101/2025.02.26.640330

**Authors:** Zahra Nozari, Paul Hüttl, Jakob Simeth, Marian Schön, James A. Hutchinson, Rainer Spang

## Abstract

**Motivation:** The cellular composition of a solid tissue can be assessed either through the physical dissociation of the tissue followed by single-cell analysis techniques or by computational deconvolution of bulk gene expression profiles. However, both approaches are prone to significant biases. Tissue dissociation often results in disproportionate cell loss, while deconvolution is hindered by biological and technological inconsistencies between the datasets it relies on.

**Results:** Using calibration datasets that include both experimentally measured and deconvolution-based cell compositions, we present a new method, Harp, which reconciles these approaches to produce more reliable deconvolution results in applications where only gene expression data is available. Both on simulated and real data, harmonizing cell reference profiles proved advantageous over competing state-of-the-art deconvolution tools, overcoming technological and biological batch effects.

**Availability and Implementation:** R package available at https://github.com/spang-lab/harp.

Code for reproducing the results of this paper is available at https://github.com/spang-lab/harplication.

## 1 Introduction

Tissues consist of cells of different types. The relative frequencies of cells of specific types define the cellular composition of a tissue, which holds crucial information on its biology and pathology. It is altered in diseases such as cancers, chronic inflammations, or infections. While cell types can be coarsely distinguished by their shape, molecular data allows for a more finely granulated distinction of cells and even cell states. The more molecules considered, the better cells can be characterized.

Cellular composition can be assessed experimentally using single cell technologies such as fluorescence-activated cell sorting (FACS; [1]), cytometry by time-of-flight (CYTOF; [2]), single-cell RNA sequencing (scRNA-seq; [3]), or combinations of these methods. However, for solid tissues, a common limitation of these approaches is the bias introduced by enzymatic dissociation, which tends to disproportionately affect certain cell types, leading to their preferential loss during isolation [4, 5, 6].

An alternative approach is bulk gene expression profiling combined with computational deconvolution [7]. In this method, a bulk expression profile is modeled as a weighted sum of reference profiles from individual cell types, where the weights represent the cellular composition of the tissue.

Let *X* be a *g*×*q* matrix representing reference profiles, where each column corresponds to a specific cell type and each row represents a gene. For the bulk data, let *Y* be a *g* × *n* matrix, where each column indicates a bulk profile and each row relates to a gene. Finally, for the cellular compositions, let *C* be a *q* ×*n* matrix where every column is a bulk tissue and every row is a cell type. The entry *C*_*ij*_ is the relative frequency of cell type *i* in tissue *j*. The central deconvolution equation connecting these data is

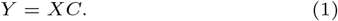

Building upon this equation, widely used tissue deconvolution tools including DTD [8], CIBERSORTx [9], MuSiC [6], or ADTD [10] estimate cellular abundances of the bulk samples. Furthermore, recent methods designed to estimate cell-type-specific gene expression, such as BayesPrism [11] and TissueResolver [12], often provide remarkably accurate cellular composition estimates as a byproduct.

Deconvolution, comes with its own limitations [13]. In theory, equation (1) should hold exactly. In reality, however, this equation does not hold, due to both tissue specific gene regulation and experimental inconsistencies in data generation. We distinguish two scenarios:

### Local Inconsistencies

*Y* = *XC* holds approximately for the majority of genes, but there is a small number of genes for which it is strongly violated. For example, if the references for T-cells were generated from inactive T-cells, while the bulk tissues contain activated T-cells. In this case equation (1) might hold for most genes, except for T-cell activation markers. Experimental inconsistencies can also lead to this problem. For example, if a certain class of genes was experimentally depleted only in the bulk profiles but not the reference profiles. In this case, equation (1) is mathematically infeasible for the depleted genes. Moreover, if reference profiles are derived from single-cell sequencing data, there can be substantial technological discrepancies compared to the bulk sequencing data used for tissues. Single-cell data is typically zero-inflated due to high drop-outs [14, 15], influenced by transcriptional burst [16], and until recently, did not commonly include ribosomal RNA depletion [17], unlike bulk RNA sequencing.

### Global Inconsistencies

*Y* = *XC* does not hold for any of the genes, because there are global inconsistencies between the bulk and reference data. This situation typically occurs if different profiling technologies such as scRNA-seq and microarrays were used [20].

Both local and global systematic differences prevent reference profiles from accurately summing up to bulk profiles. For example, Figure 1 compares bulk RNA-seq data to a weighted average of sorted RNA-seq data, with the weights determined experimentally using flow cytometry. In the UMAP plot, the measured bulk profiles and the reconstructed profiles are clearly separated.

**Figure. 1.**
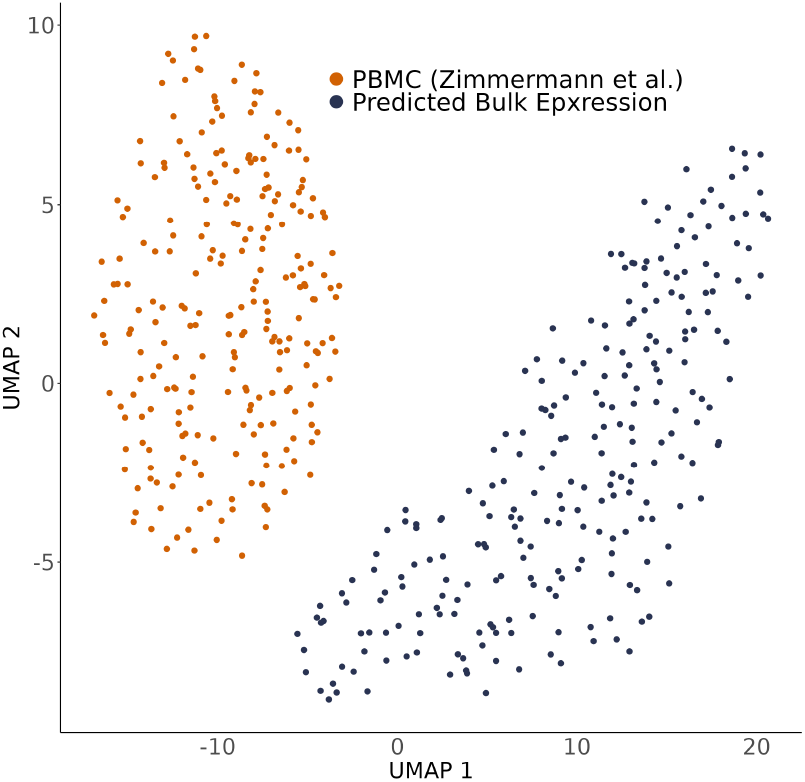
Comparison of bulk RNA-seq expression profiles to reconstructed bulk gene expression samples. Bulk RNA-seq data from [18] is represented in navy, while reconstructed bulk expression, derived by combining flow cytometry data of the same bulk samples with cell type-specific signatures from sorted RNA-seq data [19], is shown in orange.

Several approaches to compensate for data inconsistencies have been applied and described in the literature. When local inconsistencies are known, affected genes can be manually excluded from the deconvolution, as in depletion protocols. If the affected genes are not known a priori, such as in tissue-specific gene regulation, machine learning-based approaches have been proposed to detect these inconsistencies from the data. For example, Digital Tissue Deconvolution (DTD) [8] assumes that subsets of genes are affected by inconsistencies and eliminates those automatically from the deconvolution using a loss function learning approach. BayesPrism [11] in contrast, targets global inconsistencies involving all genes by marginalization of a posterior distribution conditioned on bulk and single cell expression data. CIBERSORTx provides two custom batch effect removal strategies. The first estimates an explained bulk expression matrix and then applies classical batch correction [21] to adjust this estimation to the actual bulk expression, which is only possible for moderate batch effects. The second approach directly adjusts the signature matrix used for deconvolution by integrating single-cell information. There, artificial bulk mixtures are generated from single-cell data and then batch corrected, using again the method in [21], in order to fit the actual bulk expression. Via non-negative least squares regression, taken into account the adjusted bulk mixtures and prior estimates of cellular frequencies, the adjusted signature matrix is then imputed. However, none of these methods combines and reconciles cellular composition vectors from single-cell analysis and deconvolution [22]. Furthermore, a method that systematically harmonizes possibly compromised cellular quantification measurements with transcriptomic data of various platforms is still lacking in the literature.

Here, we introduce Harp, a novel approach designed to resolve both biological and technological inconsistencies, and ensuring improved concordance between experimentally determined tissue compositions, deconvolution outcomes, and reconstructed bulk profiles, thereby enabling more accurate tissue deconvolution.

## Method

### 2.1 Algorithm

An overview of the Harp framework is provided in Figure 2. Harp operates in two modes: *Training* and *Deconvolution*.

**Figure. 2.**
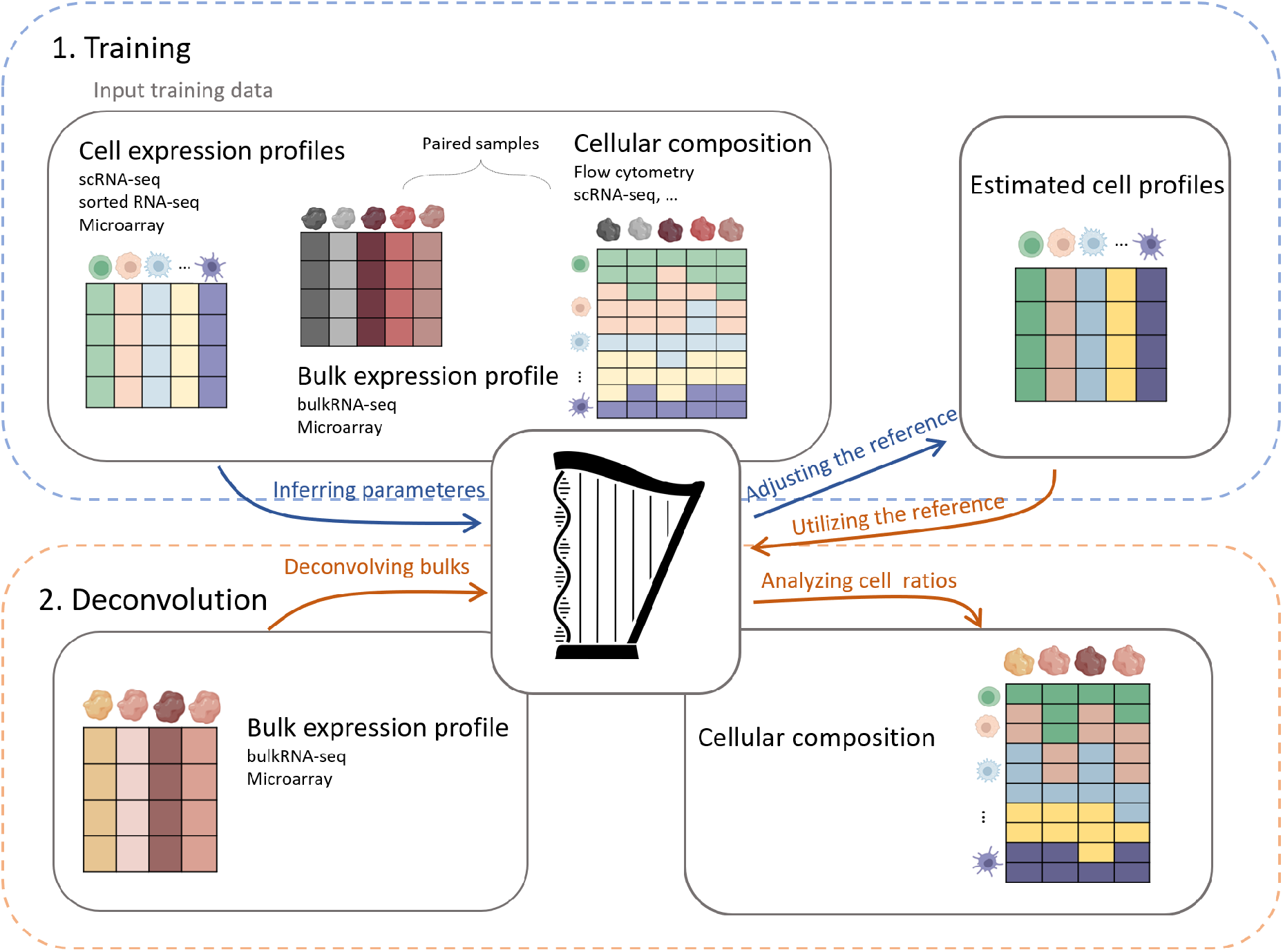
The Harp framework consists of two main modes: *Training* and *Deconvolution*. During *Training*, both bulk expression data and experimentally determined cell frequencies are used, whereas *Deconvolution* relies solely on expression profiles. In the *Training* step, Harp takes the following inputs: (1) a matrix of reference cell profiles derived from experiments on sorted cell populations using either RNA-seq or microarrays, or from single-cell RNA sequencing (scRNA-seq); (2) bulk gene expression profiles obtained from either bulk RNA-seq or microarray technology; and (3) a corresponding cellular composition matrix, generated using methods such as scRNA-seq, flow cytometry, or other techniques. Using these input data, Harp estimates a matrix of harmonized cell reference profiles. In the *Deconvolution* step, Harp takes new bulk gene expression samples, along with the estimated reference profiles from the *Training* step, to infer cellular compositions.

#### Training Mode

In this mode, Harp takes as input a matrix *Y* of bulk tissue gene expression profiles, a corresponding matrix *C* of experimentally determined cellular tissue compositions, and a matrix of reference profiles *X*, holding cell signatures of these tissues. These inputs may exhibit inconsistencies such that *Y* ≠ *XC*. The aim of Harp is to harmonize these inputs by adjusting both *X* and *C* to meet the following objectives:

- The adjusted cellular compositions, *C*^′^, accurately represent the cellular composition of the tissue.
- The adjusted reference profiles, *X*^′^, reflect the expression states of the cell types as they exist in the tissue.
- The relationship *Y* ≈ *X*^′^*C*^′^ is approximately satisfied, where *X*^′^ and *C*^′^ are the harmonized versions of the input reference profiles and cellular compositions, respectively.

First, Harp accounts for potential errors in the composition matrix *C* by allowing some flexibility, facilitating data harmonization. In order to correct for these errors, it represents the cellular decomposition of the tissue by a parameterized matrix *C*^′^(***α***) = *diag*(***α***)*C*^*^, where

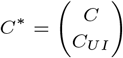

is a (*q* + 1) × *n* matrix that is identical to *C* but includes an additional row representing a mixture of all cell types that could be present in the tissue but are not accounted for in the cellular composition matrix [23, 10](see section A.1.3 for more details). The diagonal matrix *diag*(***α***), where ***α*** is a (*q* + 1)-dimensional vector of non-negative scaling parameters accounts for cell-type-specific losses during the experimental determination of *C*. Then, since cell reference data often contains signatures from more cell types than those measured in flow cytometry data, *C*, we introduce an additional column to *X*, denoted as *X*_*UI*_. This column represents the gene-wise average expression across all remaining cell types not captured in the flow cytometry data. We define this reference matrix as *X*^*^ = (*X, X*_*UI*_).

Next, Harp adjusts the reference matrix *X*^*^ to *X*^′^ to meet two key criteria:

a. a)*Y* ≈ *X*’ *C*’ (*α*).
b. b) The columns of *X*^′^ maintain similarity to the original anchor matrix *X*^*^.

Criterion b) is crucial to avoid artifacts caused by underdetermination, where *Y* = *X*^′^*C*^′^(***α***) may hold perfectly, but the adjusted profiles in *X*^′^ no longer reflect the true biological expression patterns of the cell types they represent. More formally, we minimize the following loss function with respect to *ϕ*^*^ and ***α*** simultaneously, where *ϕ*^*^ = (*ϕ, ϕ*_*UI*_) with *ϕ* being a matrix of same dimension as *X* and *ϕ*_*UI*_ being an additional column accounting for the Unidentified cell types in compatibility with the extra row in *C*^*^,

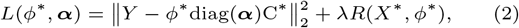

with ∥ · ∥_2_ denoting the Frobenius norm and *R* being defined as

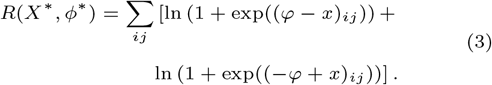

Note that the regularization term *R* constrains the adjusted references *X*^′^ to remain close to the measured references. It anchors *X*^′^.

Starting with diag(***α***) being equal to the identity matrix, the optimization process alternates between updates of *X*^′^ using

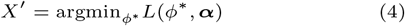

followed by the update of ***α***, by minimizing the loss function

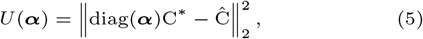

with respect to ***α***, where *Ĉ* is the estimated cellular composition using *X*^′^ as the reference profile, for more details see supplemental section A.1.5. Note that after *X*^′^ is updated, its columns are normalized to sum up to the number of features (genes) it includes; this is necessary to keep both *X*^′^ and ***α*** identifiable.

Additionally, we provided the option for Harp to automatically determine the regularization strength from a given range of *λ* values via a cross validation approach in order to arrive at an optimal *λ*^′^ that balances both criteria a) and b), see supplemental Section A.1.2.

#### Deconvolution Mode

In this mode, Harp uses the adjusted reference profile matrix *X*^′^, obtained during training, to deconvolve bulk tissue gene expression data sources similar to those used in training (e.g., comparable tissues or profiling technologies) where no experimentally determined tissue composition is available. By default, Harp applies the adjusted matrix *X*^′^ in combination with the DTD algorithm [8] for deconvolution in order to arrive at its final cell proportion estimate *C*^′^, see also supplemental Section A.1.5. However, the harmonized matrix *X*^′^ can also serve as a reference profile for use with other deconvolution methods.

### 2.2 Performance metrics

Here, we introduce several established performance metrics to evaluate deconvolution results. They either compare cell abundance estimates to ground-truth cell proportions—which may either be predefined in simulation scenarios or obtained experimentally (e.g., through flow cytometry or scRNA-seq)—or predicted bulk gene expression to observerd bulk expression data. For their mathematical definition we refer to supplemental Section A.1.4.

#### Cell Type-Specific Performance

The first performance metric of a deconvolution tool, *R*_*c*_(*l*), is its ability to accurately capture variations in the relative abundance of a given cell type *l* across different samples. For example, this quality metric can evaluate how accurately the method predicts that sample 1 contains 10% more T cells than sample 2.

#### Sample-Specific Performance

A second performance metric is the tool’s ability to accurately estimate the proportions of different cell types within an individual sample *m*, denoted as *R*_*s*_(*m*). For instance, *R*_*s*_ can evaluate how accurately the method determines that one sample contains 30% more T-cells than B-cells.

#### Combined Performances

Following [6] the cell type-specific and sample-specific performance metrics can be integrated into a single overall measure denoted as R. We furthermore include the absolute quality metrics RMSD and mAD into our analyses.

#### Bulk Reconstruction Performance

Thus far, our performance metrics have focused on evaluating the estimated cellular compositions *C*. In addition to accurately recovering these compositions, a well-calibrated deconvolution tool should also be able to recapitulate the observed bulk expression *Y* via the model

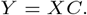

To assess this, we introduce a bulk-centered performance measure *ρ*(*m*), which correlates explained bulk expression of sample *m* to its observed bulk expression.

## 3 Simulation

To evaluate the performance of Harp and compare it with state-of-the-art deconvolution algorithms, we performed extensive benchmarking simulations. In these simulations, we followed a well-established approach by generating artificial bulk expression profiles as weighted averages of single-cell expression profiles [9, 6, 11]. Importantly, the simulated profiles allow us to control the “cellular composition” through the assignment of weights. For instance, in a profile where T cells constitute 30% of the cellular composition, the cumulative weight of the T cell profiles accounts for 30% of the total. In our simulations, we used single-cell data from two studies on non-Hodgkin lymphomas (nHL) [24, 25]. The study by Steen et al. [24] comprises profiles from 28,416 single cells collected from eight patients, including four patients with Diffuse Large B-cell Lymphoma (DLBCL), three patients with Follicular Lymphoma (FL) and one control patient with Tonsilitis (T). The cells have been pre-annotated with the following cell type labels: B cells, Monocytes, Natural Killer cells, Plasmablasts, CD4 T cells, CD8+ T cells, regulatory T cells, T follicular helper cells, and a remaining unknown compartment. The second study, conducted by Roider et al. [25], includes 35,284 single-cell profiles from 12 nHL cases, including three DLBCL, four FL, two Transformed Follicular Lymphoma (tFL) and three control patients exhibiting reactive lymph nodes (rLN). Similarly, these cells have been pre-annotated as B cells, myeloids, CD8+ T cells, regulatory T cells, follicular helper T cells, and T helper 1 cells. Notably within this study the B cells were further divided into healthy B cells and malignant lymphoma cells. For further details see supplement, Section A.2.2 and Figure 9.

We observed substantial inconsistencies and batch effects between these studies (also see supplemental Figure 8a), which can be partially attributed to differences in laboratory protocols, tissue handling, and library preparation, as well as patient heterogeneity (see supplemental material, Section A.2.1). We used these discrepancies to simulate inconsistent data sets. Specifically, the data from the first study defined the reference profiles, while the data from the second study were used to generate artificial bulk samples. For Harp’s anchor *X*^*^ we used the average cell-type specific expression across single cell profiles of [24]. Concerning [25], we randomly divided the data from 12 patients into two independent sets of six patients each. One set was used to generate a training set of artificial bulk samples, while the other set was used to create an independent test set. For deconvolution we focused on only those cell types that were defined in both studies, namely B cells, CD8+ T cells, regulatory T cells and follicular helper T cells. Importantly, the B cell compartment contained both malignant and physiological cells from this B cell malignancy. This implies completeness of the reference and thus, *X*^*^ = *X*.

For each bulk sample, we randomly selected cells stemming from a single patient only. More precisely, we determined the actual amount of single cells for each cell type within a given patient and then perturbed this amount with a normally distributed factor in order to arrive at the quantity of cells to be randomly selected from each cell type. This allowed us to generate multiple artificial bulk mixtures from a single patient, which contain suitable variation in terms of cellular compositions, see supplemental Section A.2.3 for details. Following [11], we introduced additional distortions that further amplify the discrepancies observed between the datasets, by locally perturbing gene expression values in artificial bulk mixtures with a gene specific multiplicative noise. More precisely, we sampled gene-specific factors from a pre-defined normal distribution and then multiplied this factor to the expression value of the considered gene in all artificial bulk mixtures. Following this approach, we perturbed 40% of all genes while the remaining set of genes was left unchanged (see supplement, Section A.2.4, for details). In total, we generated 50 artificial training samples and 40 test samples using this protocol (also see supplement, Section A.2.5). Supplemental Figure 11 shows the distribution of cell proportions in these datasets. Importantly, this simulation framework naturally controls the proportions of the various cell compartments in the artificial bulk mixtures, see supplemental Section A.2.3 for details.

### 3.1 Calibration of regularization

Our first analysis addresses the calibration of the parameter *λ* in equation (2). Regularization enforces a degree of similarity between the adjusted reference matrix *X*^′^ and its unadjusted counterpart *X*^*^. Note that overly strong regularization results in minimal adjustment of the reference profiles, potentially failing to compensate for technical discrepancies. On the other hand, weak or absent regularization can produce reference profiles that diverge from the true expression characteristics of the cells they represent. For example, the column corresponding to B cells in *X*^′^ might no longer capture the typical expression profile of a B cell, indicating that the reference has been “over-adjusted”.

In order to better understand Harp’s dependency on its hyperparameter *λ*, we fitted models on simulated data using different *λ* values in the range [0, 2^15^]. Let *X*^′^(*λ*) be the adjusted reference matrix produced by Harp when using regularization strength *λ*. Figure 3 shows a UMAP [26] embedding of (a) the single-cell profiles used in the training data and (b) the columns of *X*^′^(*λ*) containing the adjusted reference profiles for various cell types at different values of *λ*. In the plot, small dots represent single-cell expression profiles, with colors indicating their corresponding cell types. In contrast, squares denote reference profiles extracted from *X*^′^(*λ*), where the square size increases with larger *λ* values. Triangles represent the reference profiles obtained for the optimal *λ*^′^, which are the adjusted references used by Harp in *Deconvolution* mode, see Section 2.1.

**Figure. 3.**
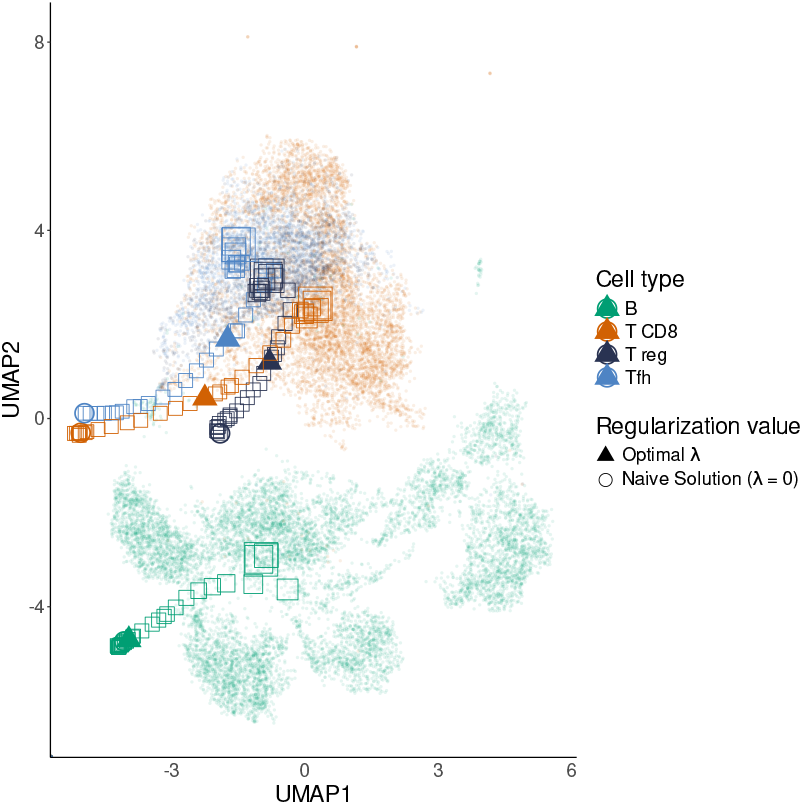
Reference profiles learned by Harp for different regularization parameter (*λ*) values, embedded in the single-cell context using UMAP. Each square represents a cell type-specific expression profile in the Harp reference for a given *λ*, with square size encoding the magnitude of *λ*. Triangles indicate the optimal estimated reference profiles selected by Harp, and each color corresponds to a specific cell type.

We observed that large *λ* values yield reference profiles located at the centers of the corresponding single-cell clusters, indicating minimal adjustment. As *λ* decreases, adjustments become visible as the reference profiles gradually shift away from the cluster centers; however, except for very small *λ* values, they remain in the vicinity of the clusters they represent. Optimal reference profiles (shown as triangles) tend to lie along this trajectory toward the center, reflecting the typical bias-variance trade-off observed in machine learning applications.

### 3.2 Benchmarking

We benchmarked Harp’s performance against a set of widely used deconvolution tools, including BayesPrism [11], CIBERSORT [27], CIBERSORTx [9] and MuSiC [6]. Harp was trained on the training set of bulk mixtures with known ground truth proportions and subsequently evaluated in *Deconvolution* mode on the independent test dataset. Since none of the other algorithms incorporate a training phase for data harmonization, they were applied directly to the bulk samples in the test data, see supplemental Section A.2.6 for details. Figure 4 demonstrates that in these simulations, Harp outperformed its competitors across all performance metrics introduced in Section 2.2. Particularly pronounced differences were observed in the cell type-specific correlation for the least abundant cell type, regulatory T cells. In this case, the competing algorithms fell well behind Harp’s performance, suggesting that integrating a training mode aiming at data harmonization was especially effective for accurately inferring proportions of rare cell types.

**Figure. 4.**
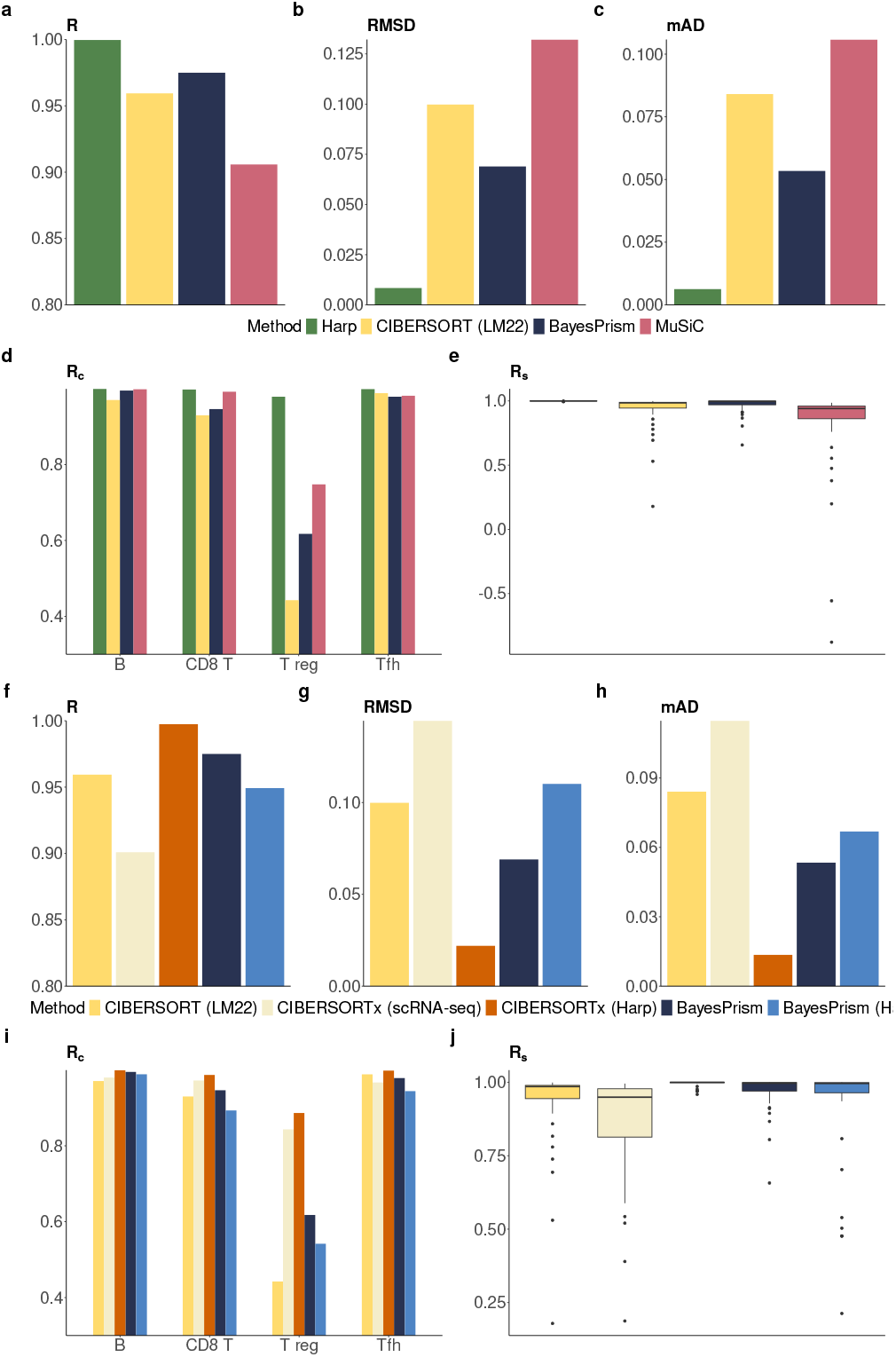
Evaluation of performance metrics in simulations. (a-e) depict the main benchmark of comparing Harp to the default of competing algorithms, whereas (f-j) depicts the hybrid deconvolution scenario of providing the harmonized reference matrix, *X*^′^, calculated by Harp to other deconvolution tools. (a-c) and (f-h) show the combined performances R, RMSD, and mAD; (d) and (i) the cell type-specific performance, *R*_*c*_, and (e) and (j) the sample specific performances. In brackets we notated the exploited reference, where LM22 is the custom reference provided by CIBERSORT and scRNA-seq refers to the internal CIBERSORTx method of constructing a custom reference from provided scRNA-seq data.

One might argue that, in the previous benchmark, Harp had an advantage, because it was trained with additional data, including ground-truth compositions, and could perform data harmonization—a capability not available to the competing algorithms. To test whether harmonization could also improve the performance of these methods, we provided the harmonized reference matrix, *X*^′^, calculated by Harp to competing deconvolution tools, see supplemental Section A.2.6 for details. We then evaluated these hybrid methods, which combine Harp’s harmonized reference *X*^′^ with the deconvolution algorithms of CIBERSORTx and BayesPrism. Figure 4 shows that the harmonized reference learned by Harp is beneficial for deconvolution with CIBERSORTx. In particular, using the Harp reference significantly improved all performance scores compared to both the default LM22 and customized scRNA-seq references. Interestingly, hybrid CIBERSORTx with the Harp reference outperformed BayesPrism, even though BayesPrism leverages full single-cell information. For BayesPrism with the Harp reference, we did not observe a general improvement; however, performance scores, including sample-specific correlations, indicate that its performance is on par with that of CIBERSORT (LM22). As observed previously, a notable strength of the Harp reference is its performance in the lowly abundant regulatory T cell compartment. In this case, using a generic reference (CIBERSORT (LM22)) or relying solely on single-cell information (BayesPrism and CIBERSORTx (scRNA-seq)) appears to miss the fine-grained cellular signals present in the bulk expression data.

### 3.3 Uncertain experimental compositions

Thus far, our evaluations have assumed that the composition matrix *C* is predetermined by the simulation design and used as ground truth. In practical applications, however, *C* is determined experimentally and may be subject to bias. Harp aims to partially correct for this bias through the cell type-specific correction rates, ***α***, incorporated into its loss function (see equation (2)). In the following simulation experiment, we evaluate the effectiveness of this correction.

We simulated training and test bulk mixtures as described above. By design, these simulations yielded the correct cell-type proportions *C*(*l*). We then additionally multiplied these proportions by a random, multiplicative, cell type-specific distortion ***δ***(*l*) (with *l* representing a cell type), introducing a bias analogous to that caused by cell loss or systematic gating errors. Although this distortion may vary across cell types, it was kept constant for all samples in the training data, see supplemental Section A.2.7 for details.

We then ran Harp in *Training* mode twice: once using the correct cell proportions and once with the distorted proportions. Figure 5 demonstrates that the distortions had minimal impact on Harp’s overall performance. Moreover, examining the estimated parameters *α*(*l*) alongside the distortion rates ***δ***(*l*) reveals that, as expected, ***α***(*l*) ≈ ***δ***(*l*)^−1^, see supplemental Figure 12.

**Figure. 5.**
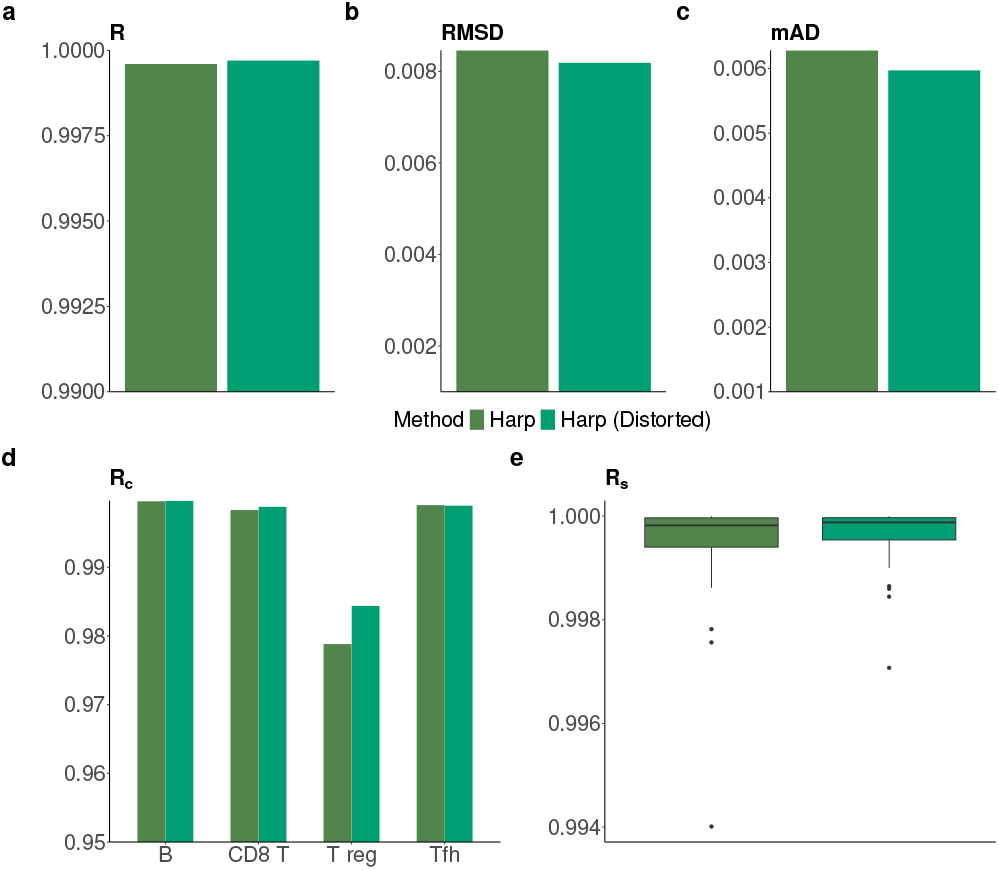
Comparison of Harp when provided with correct cellular proportions vs. distorted proportions. The distorted case is indicated by (Distorted). (a-c) show the combined performances R, RMSD and mAD; (d) represents the cell type-specific performance, *R*_*c*_; and (e) depicts the sample specific performances, *R*_*s*_.

## 4 Data harmonization with Harp improved deconvolution accuracy in a study combining data from two distinct sources

We assessed Harp’s performance by integrating data from two sources. The bulk RNA-seq data were obtained from a study investigating primary peripheral blood mononuclear cells (PBMCs) in healthy individuals following influenza vaccination [18]. We utilized bulk RNA-seq expression profiles for 250 cases, with the cellular composition of the PBMCs experimentally determined via flow cytometry. However, this study did not include reference profiles for the various PBMC cell types. For the cell expression references we used data from an independent source [19], which generated RNA-seq profiles of sorted PBMC cell populations. Details on data prerocessing can be found in supplemental Section A.3.1.

We split the bulk data into a training set of 150 cases and a test set of 100 cases. For the training set, we ran Harp using the reference data from the second source as the anchor *X*^*^ for regularizing the reference profile. We then applied Harp in *Deconvolution* mode to the test bulk samples, alongside CIBERSORT and BayesPrism, and compared the deconvolution results to the corresponding flow cytometry measurements (see details on the configurations of other algorithms in section A.3.2).

Figures 6 (a-g) show that Harp outperformed its competitors in several, though not all, performance metrics. Most notably, it achieved robust overall performance, as indicated by the metrics R, RMSD and mAD, which were supported by excellent sample-specific reconstructions of cell proportions.

**Figure. 6.**
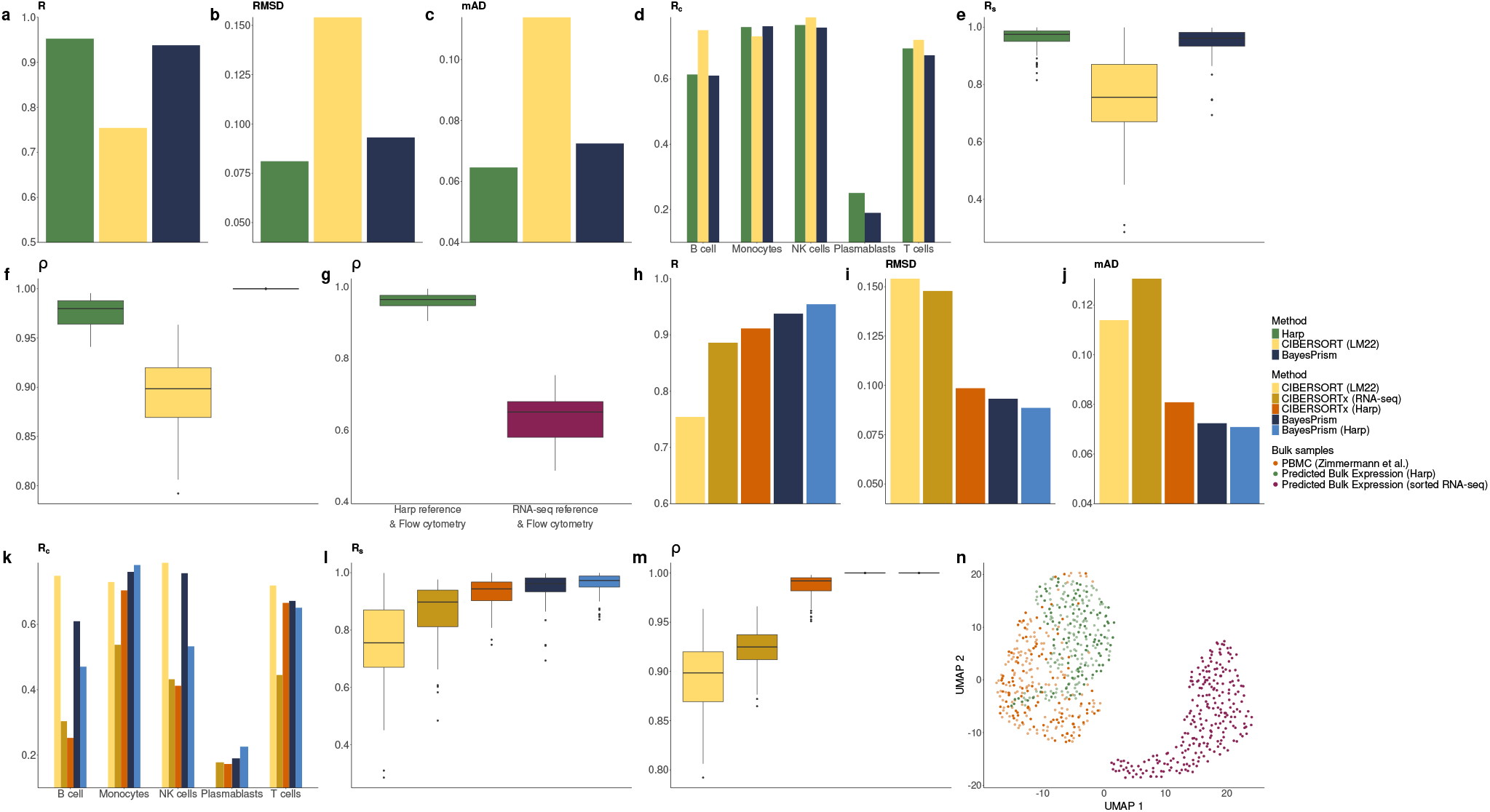
Evaluation of performance metrics in the deconvolution of 100 PBMC RNA-seq test samples with a sorted RNA-seq reference. (a-f) represent the benchmark of deconvolution tools (Harp, CIBERSORT (LM22), and BayesPrism). Plots (a-e) evaluate performance on the prediction of cell proportions, while plot (f) analyses the quality of the reconstructed bulk gene expression profiles. (g) shows box plots of Pearson correlations *ρ* between the reconstructed bulk gene expression profiles, using the experimental cellular compositions in conjunction with the Harp (green) and sorted RNA-seq derived (magenta) references, respectively, and the observed bulk RNA-seq data. (h-m) depict the hybrid deconvolution scenario of providing the harmonized reference matrix, *X*^′^ (sorted RNA-seq reference as the anchor *X*^*^), calculated by Harp to CIBERSORTx and BayesPrism, and these results are compared to those from the respective default methods. Plots (h-l) assess the accuracy of the estimated cell proportions, while plot (m) evaluates the quality of reconstructed bulk gene expression profiles. (n) is a UMAP of the predicted bulk gene expressions of 100 PBMC test samples (darker color shades) and 150 PBMC training samples (represented in lighter color shades), using the Harp (green) and sorted RNA-seq (magenta) reference, respectively, in conjunction with the cellular compositions derived from experimental data. This plot also includes observed bulk RNA-seq expression profiles (orange).

In addition to this evaluation of cell proportion reconstruction, we evaluated data consistency after harmonization by reconstructing bulk expression profiles *Y* using reference data *X* and cell compositions *C* according to the formula *Y* = *XC*. We performed this reconstruction twice: once using the original (anchor) reference *X*^*^ and once using the harmonized reference *X*^′^ estimated by Harp. The cell abundances, *C*, used in both sets of reconstructed bulk samples were obtained from flow cytometry data (also see section A.3.3). Figure 6 (g) shows that the bulk reconstructions based on Harp’s reference exhibited a higher correlation with the observed bulk profiles. This improved consistency was even more evident when both observed and reconstructed bulk profiles were embedded in a UMAP, see Figure 6 (n).

We also compared the performance of the methods in terms of the reconstruction of bulk samples using each method’s corresponding reference and estimated cell proportions (see details in supplemental Section A.3.2). As shown in Figure 6.f, Harp performed better than CIBERSORT(LM22), while BayesPrism achieved the best performance. However, in regard of this quality metric BayesPrism always showed the strongest correlation (≈ 1) independent of the provided dataset. This is likely due to an explicit constraint in the method’s optimization approach, which forces reconstructed expressions to match the original data values [11].

Similarly to the simulation experiments, we provided the harmonized reference matrix learned by Harp to CIBERSORTx and BayesPrism, and compared the performance of this approach, to that achieved with their default LM22 reference (see supplemental Section A.3.2 for details). Figures 6 (h-j), show that Harp’s reference improved the overall performance of both methods. Moreover, cell proportions within samples were better reconstructed when using the harmonized references (Figure 6 (l)). For predicting cell type-specific abundance, *R*_*c*_, see Figure 6 (k), CIBERSORTx benefited more from using Harp’s reference than from constructing a custom reference from sorted RNA-seq data, although CIBERSORT (LM22) performed even better. Additionally, we observed that the Harp reference improved BayesPrism’s prediction of monocyte and plasmablast proportions, while it had a negative impact on the predictions for B cells, NK cells, and T cells proportions. Regarding the prediction of bulk gene expression (see Figure 6 (m)), BayesPrism outperformed the other methods in both its default mode and when using the Harp reference. Moreover, with the harmonized Harp reference, CIBERSORTx showed improved performance compared to both CIBERSORTx (RNA-seq) and CIBERSORT (LM22).

In summary, we observed clear gains in overall and sample-wise performance, as well as bulk reconstruction ability when using Harp’s reference compared to a method’s default reference, though cell type-specific performance yielded mixed results.

So far we used data from different sources but comparable technologies, as both the bulk profiles and the anchor reference samples were derived using standard RNA-seq protocols. Next we challenged Harp and its competitors further by using microarray-derived reference profiles as a starting point (anchor *X*^*^) for harmonization. Our analysis is identical to that described above, with the sole difference that CIBERSORT’s LM22 matrix, which is microarray-derived, replaces the references derived from RNA-seq profiles of sorted cell compartments (for details, see supplemental Section A.3.2). Figures 7 (a-e) show that, for cell proportion predictions, Harp outperformed both methods across all performance metrics, except for *R*_*c*_ in the B cell compartment. Nonetheless, the performance of all methods with respect to the *R*_*c*_ metric was comparable for most cell types. Figure 7 (f) shows that both Harp and BayesPrism, when using LM22, explained the bulk gene expression samples better than CIBERSORT. Moreover, as shown in Figure 7 (g), the advantage of using the Harp estimated reference over the microarray-based LM22 reference for bulk reconstruction was pronounced, with the average correlation improving from approximately 0.5 to about 0.9 (see also supplemental Section A.3.3).

**Figure. 7.**
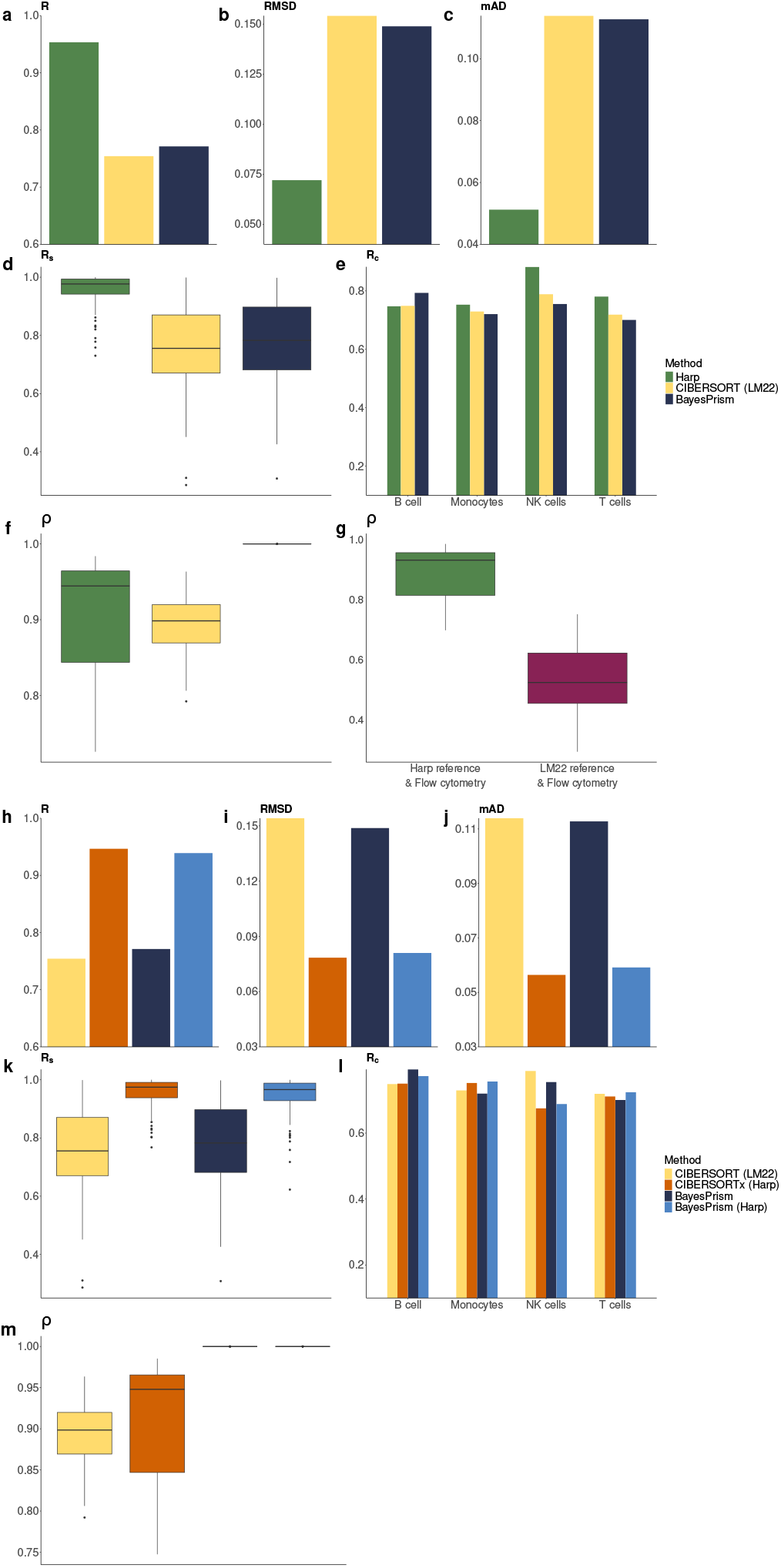
Evaluation of performance metrics in the deconvolution of 100 PBMC RNA-seq test samples with a microarray-based reference (LM22). (a-f) represent the benchmark of various deconvolution tools (Harp, CIBERSORT (LM22), BayesPrism). Plots (a-e) evaluate performance in predicting cell proportions, while Plot (f) analyses the quality of the reconstructed bulk gene expression profiles.(g) shows box plots of the Pearson correlation, *ρ*, between the reconstructed bulk gene expression profiles, using the flow cytometry-derived cell proportions, together with the Harp(green) and LM22 (magenta) references. (h-m) show the hybrid deconvolution scenario of providing the harmonized reference matrix, *X*^′^ (microarray-based reference, LM22, as the anchor *X*^*^), calculated by Harp to CIBERSORTx and BayesPrism, and the results are compared to those from their respective methods. Plots (h-l) evaluate performance in predicting cell proportions, while plot (m) analyses the quality of the reconstructed bulk gene expression predictions.

**Figure. 8.**
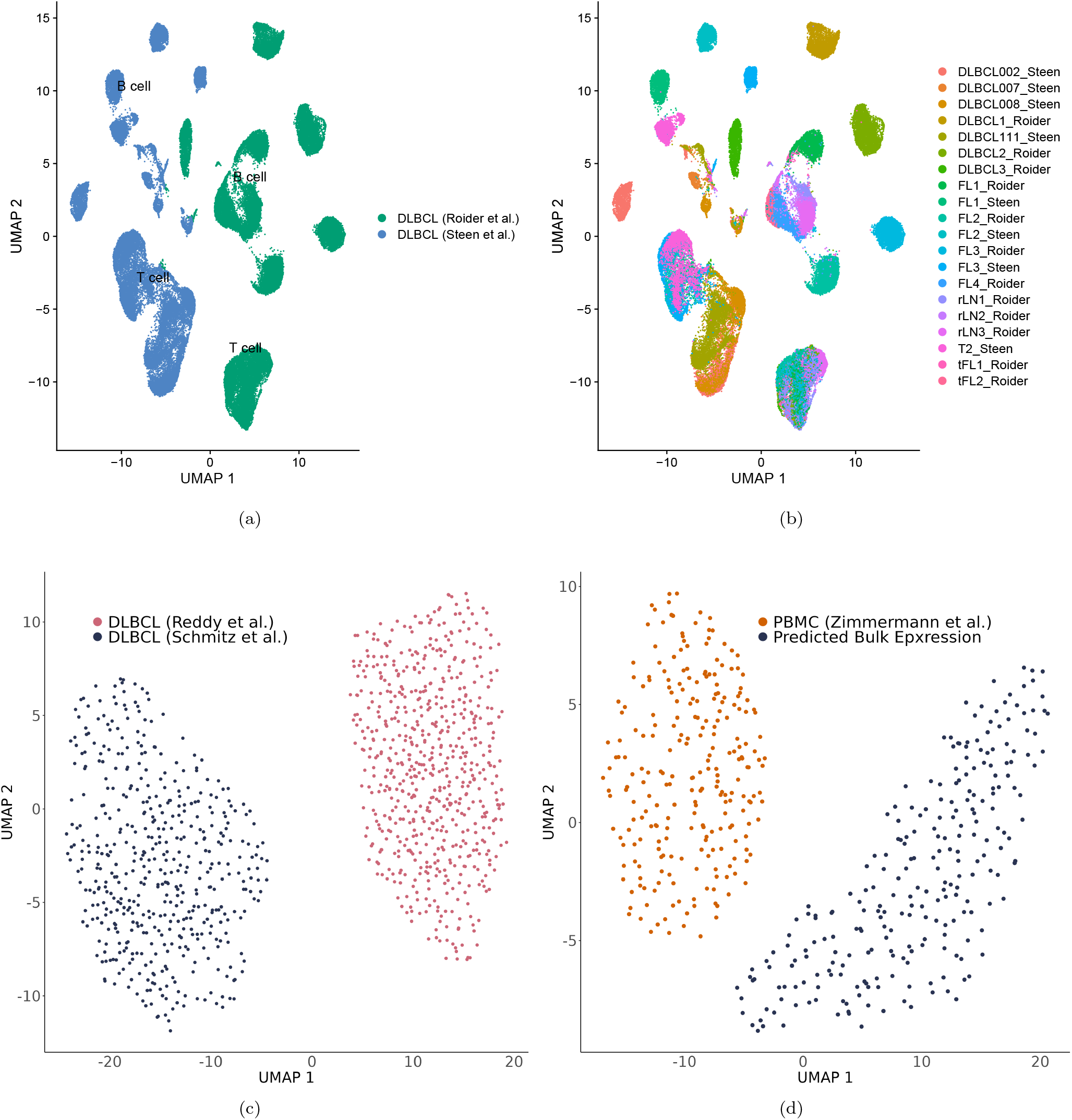
(a) DLBCL single-cell profiles from two distinct studies: green represents data from Roider et al. [25], while blue corresponds to Steen et al. [24]. (b) Same data as in (a), but colored by patient sample from the respective study. (c) DLBCL bulk RNA-seq expression data from two studies: pink refers to data from Reddy et al. [28], and dark blue illustrates Schmitz et al. [29] data. (d) Bulk RNA-seq expression of samples from [18] is represented in orange, while reconstructed bulk expression—derived by multiplying flow cytometry data of the same bulk samples with cell signatures from sorted RNA-seq data [19]— is shown in navy.

**Figure. 9.**
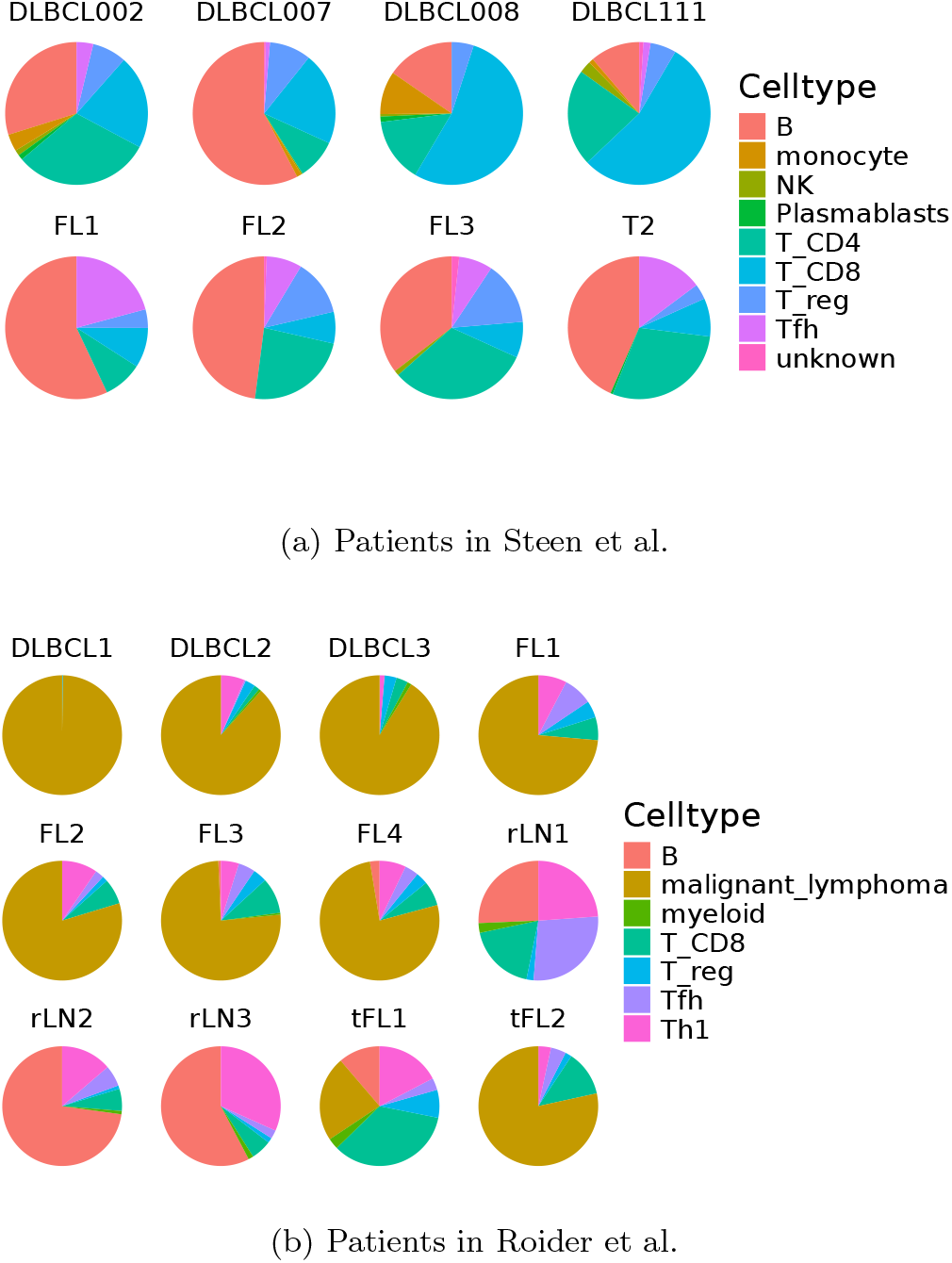
Cellular composition of the patients from Steen et al. and Roider et al. The lymphoma entities comprise Diffuse Large B-cell Lymphoma (DLBCL), Follicular Lymphoma (FL), Transformed Follicular Lymphoma (tFL), Tonsilitis (T) and Reactive lymph nodes (rLN)

Finally, we again examined the effect of using harmonized references on the performance of competing tools. Figures 7 (h-k) and Figure 7 (m) show that both CIBERSORT and BayesPrism benefited from using Harp’s reference. Notably, when CIBERSORTx was used with Harp’s reference, no batch correction was performed when applying CIBERSORTx, yet its performance still improved. Regarding the prediction of cell type-specific proportions, Figure 7 (l) indicates that both methods achieved comparable performance across different references, with the exception of NK cells, where performance was higher when using each method’s default reference.

The benchmark comparison of deconvolution tools and Hybrid deconvolution on microarray bulk expression data, where technological inconsistency is insignificant, is discussed in Supplemental Section A.4. Therefore, in this case, harmonization was not particularly required. The results indicate that Harp’s performance is comparable to its performance discussed earlier.

## 5 Discussion

We introduced Harp, a novel deconvolution tool designed for applications where reference data and bulk data are derived from different sources and are therefore not fully compatible. By performing data harmonization, Harp overcomes these discrepancies, emerging as a cross-platform deconvolution tool that enables analyses beyond the confines of a single data source.

We emphasize that in our work harmonization is not an end in itself; the ultimate goal is to accurately predict the cellular composition of a tissue. In its *Training* mode, Harp uses experimentally determined proportions of various cell compartments, adapting the reference profiles to achieve better overall consistency between these proportions, the reference data, and the bulk expression. However, the input cell compositions may be compromised by cell loss during tissue preparation, the omission of cell compartments that were present in the tissue, or errors during gating (manual or automated). As a result, pushing deconvolution results closer to these potentially flawed experimental measurements—as Harp does in *Training* mode—might be counterproductive, even if the deconvolution outcomes appear to better match the experimental data, as observed in our evaluations.

Moreover, deconvolution tools should not be seen merely as a way to replicate flow cytometry analyses. It is possible that computational deconvolution, in some cases, could yield more accurate estimates than experimental quantifications, as it accounts for signals from all cell compartments within the tissue—potentially capturing components that might otherwise be overlooked.

However, this leaves us with the challenge of determining which approach is more accurate, as a definitive ground truth does not currently exist [13]. We anticipate that this will change as both experimental protocols and image-based analyses progress rapidly. In the meantime, we advocate harmonizing all available information so that apparent discrepancies (e.g., those shown in Figure 1) are addressed. Harp is designed to achieve precisely this.

## Acknowledgments

This project has received funding from the European Union’s Horizon 2020 research and innovation program under the Marie Sk-lodowska-Curie grant agreement No 860003 as well as Bundesministerium für Bildung und Forschung (BMBF, German Federal Ministry of Education and Research) [031L0173].

## Appendix

### A.1 Method

#### A.1.1 Optimization algorithm

We numerically solved the optimization problem described in Section 2.1 via a custom gradient descent approach. We chose the anchor reference profile *X* as initial value for *ϕ* in the gradient descent. This is motivated by the fact that the final optimal reference profile *X*^′^ shall encode biological information by perturbing *X* along the optimization process.

The partial derivative of the loss function, see equation (2), can be written as

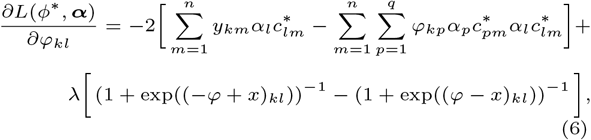

where *φ*_*kl*_ is the expression level of gene *k* in cell type *l*. Using this gradient, we evolve *φ* via gradient descent

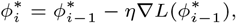

where *η >* 0 denotes the learning rate.

In order to automatically determine an optimal learning rate in each step, especially for balancing runtime efficiency and accuracy, we used Armijo back tracking, see [30]. More precisely in each gradient descent step the learning rate is iteratively decaied with the factor *γ*^*k*^ where *γ* ∈ (0, 1) and *k* = 0, 1, 2 … until the sufficient decrease condition

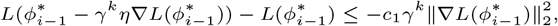

is met. In our applications we chose *c*_1_ = 10^−4^ and *γ* = 0.5. As convergence criterion we required smallness of the gradient, i.e., we stopped when

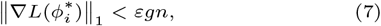

where we chose the hyperparameter *ε* = 10^−4^. Due to the dimensionality of the problem we included both the number of genes, *g*, and the number of samples, *n*, in the breaking condition. However, in order to guarantee robust breaking in the case of non-convergence we limited the number of iterations to 10^5^ and the minimal *η* during backtracking to 10^−7^. We note, however, that in our applications, we always satisfied condition (7).

##### A.1.2 Regularization approach

In the loss function *R* (see equation (2)) serves as a regularization term controlled via the parameter *λ*. This regularization is essential to ensure the biological integrity of the reference profiles. Practically, it ensures the incorporation of single-cell information into the derived reference profile for *λ >* 0.

The user can either provide a desired regularization value directly or Harp can determine an optimal regularization value *λ*^′^ via the following cross-validation approach. First, Harp uses a range of candidate *λ* values, which by default is given as the sequence

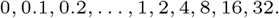

Next the provided training data set is split into *n* folds, with *n* = 5 by default. Now for each regularization value in the provided range, Harp is executed in *Training* mode with the exploited samples being comprised of *n* − 1 folds, yielding a reference *X*^′^(*λ*). With *Y* being given as the bulk samples from the remaining fold, we solve

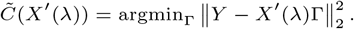

The solution is given as

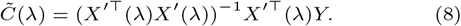

In the next step, for each cell type we calculate the correlation, *R*_*c*_, between these predicted cell abundances, 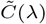, and the cell proportions from the experiments. In cross-validation, this provides us with a cell type-specific quality score for all samples in the original training set. Thus, we consider the mean correlation across all cell-types and samples in order to arrive at a single score for each candidate *λ* value. The *λ* which provides the highest score is then selected as optimal *λ*^′^ implying the use of *X*^′^(*λ*^′^) for the subsequent steps of the algorithm.

The regularization term *R* in equation (3) is given via the SoftPlus function which serves the purpose of being analytic at zero while being a suitable approximation of ∥*ϕ*^*^ − *X*^*^∥_1_. The scalar SoftPlus function is defined as

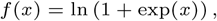

and its derivative is represented by

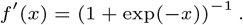

Therefore,

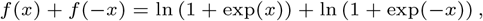

which can be approximated as

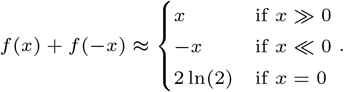

Thus, the absolute value function can be approximated as

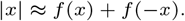

##### A.1.3 Unidentified Cell types

Reference matrices typically do not cover all cells in a tissue. This could be due to the technical effects or experimenters’ decisions depending on the set of cell types that they are interested in. Therefore, we introduce a row in *C* prior to deconvolution, which leads to estimating an additional column in *X*^′^ that represents the missing cells. The row *C*_*UI*_ that we add to the cellular composition matrix, *C*, to create

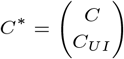

is obtained as

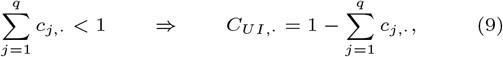

see also [23, 10].

##### A.1.4 Performance metrics

Let *C*^True^ be a *q* × *n* matrix containing ground-truth cellular proportions. Mathematically, the cell type-specific performance of a deconvolution tool as introduced in Section 2.2 is defined as

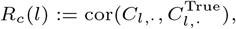

where *l* denotes a specific cell type and the correlation is calculated over all samples using only the proportions for that cell type. The sample-specific performance is defined as

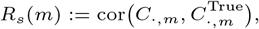

where *m* represents a specific sample.

In analogy to [6] we combine cell type-specific and sample-specific performance into a single correlation score denoted as R. To do so, we first concatenate the cell type proportions from all cell types into single vectors:

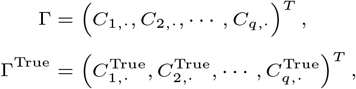

where *q* denotes the number of cell types. The combined performance is then quantified by the correlation between G and G^True^:

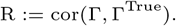

Additionally, we analyze two absolute error metrics, in analogy to [6]. The overall root mean squared distance (RMSD) is given by

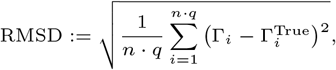

and the overall mean absolute deviation (mAD) is defined as

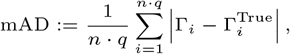

where *n* represents the number of samples.

Complementing the previous cell abundance-centric quality scores we also introduce the bulk-centric quality score

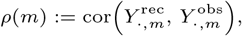

where *m* is a sample and *Y* ^obs^ denotes the observed gene expression matrix, measured through experiment, and *Y* ^rec^ is the reconstructed matrix obtained using a reference matrix *X* and a composition matrix *C* according to

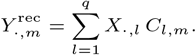

For Harp, the reconstruction *Y* ^rec^ is performed using *X*^′^ and *C*^′^, while for competing tools, the corresponding reference matrices and estimated composition matrices are used.

##### A.1.5 Digital Tissue Deconvolution (DTD)

To infer the cellular compositions *Ĉ* during Harp’s *Training* mode (see equation (5)), we use DTD [8]. During training of DTD, genes are weighted differently to improve the accuracy of the cell abundance estimation. This method consists of two nested objective functions: an outer function *L*(***g, α***) and an inner function *ℒ* _***g***_.

For a given vector ***g*** of length *g*, DTD determines *Ĉ*(***g***) via

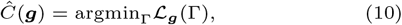

where

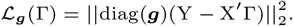

Here *Y* is comprised by the bulk samples contained in the training set. The inner function

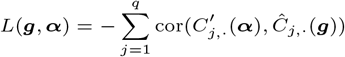

evaluates discrepancies between the cellular frequencies *Ĉ*(***α***) estimated in equation (10) and the cellular frequencies *C*^′^(***α***) = diag(***α***)C^*^ given by rescaling the experimental measurements *C*, with the factors ***α*** being given from the previous step of the iterative updating scheme of Harp’s *Training* mode, see Section 2.1. Thus, given ***α*** we solve

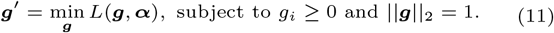

Note that with ***g***^′^, *β****g***^′^ is also a minimum of *L* for any *β >* 0. Therefore, the constraint ∥ ***g*** ∥_2_ = 1 in equation (11) is necessary to ensure uniqueness.

The minimum of L_***g***_ is given analytically via

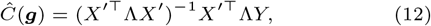

where Λ := diag(***g***). Inserting this term into *L* results in a single optimization problem in ***g***, which is minimized using a gradient descent algorithm.

Let 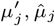 and 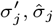 denote the mean and standard deviation of 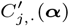 and *Ĉ*_*j*,·_ (***g***), respectively. The gradient is then computed as

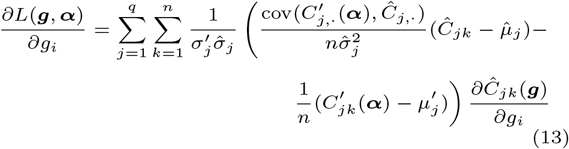

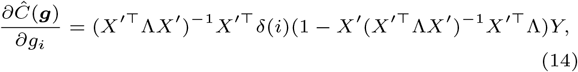

where *δ*(*i*) ∈ ℝ^*p*×*p*^ is defined as

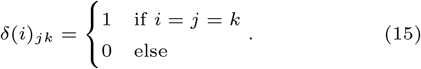

We emphasize that in the *Training* mode of Harp, DTD is trained on the real data (bulk gene expresssion and cell compositions).

In *Deconvolution* mode, Harp uses the adjusted reference profile matrix *X*^′^, obtained during training, to deconvolve bulk tissue gene expression data from similar sources (e.g., comparable tissues or profiling technologies) where no experimentally determined tissue composition is available. By default, Harp exploits the adjusted matrix *X*^′^ in combination with the scaling factor diag(***g***) determined in equation (11) in order to solve equation (10), where now *Y* represents the new bulk samples that need to be deconvolved. For applying DTD in Harp, we used the default setting as advised in the corresponding documentation.

However, the harmonized matrix *X*^′^ can also serve as a reference profile for use with other deconvolution methods.A.1

#### A.2 Simulation

##### A.2.1 Inter study inconsistencies

As described in Section 3 we used the scRNA-seq data from two studies [24, 25] as the sources for our simulations. Notably, we observed various inconsistencies between these studies arising both from technical batch effects and biological variability.

Firstly, these two datasets were processed through different library preparations. The single cell library from Roider et al., [25] was prepared by the Chromium single cell v2 3’kit (10x Genomics). Whereas Steen et al., [24] employed the 10x Chromium 5’ kit for library preparation. The two assays capture different transcript ends but use the same polydT primer for reverse transcription, see [31]. Although they have similar results in cell-type identification, there are some differences in the expression of a certain group of genes [31]. The variations in sample preparation protocols between the two studies also cause visible discrepancies between the datasets, see Figure 8a.

Secondly, as expected, we observed patient-specific inter cell-type heterogeneity especially in the malignant B cell compartment, naturally arising from patient-specific mutations in DLBCL cancer, see Figure 8b. We stress that this patient heterogeneity carries essential biological information on the one hand; however, it also makes naive, coarse-grained deconvolution impossible. Thus, preserving this variability in our simulations challenged deconvolution tools to account for patient heterogeneity when predicting cellular composition. In conclusion, by generating artificial bulk mixtures from [25] and using [24] as single-cell library in simulations, we naturally accounted for various sources of heterogeneity yielding inconsistencies between bulk mixtures and single-cell references that needed to be accounted for during deconvolution.

##### A.2.2 Preprocessing of single cell data for simulations

The downstream preprocessing of the single cell libraries [24] and [25] was performed exactly as in [32]^1^, where both datasets were analyzed for their most variable genes, arriving at the joint 1000 most variable genes.

##### A.2.3 Bulk oversampling

The following oversampling step was included in the generation of pseudo-bulks in order to arrive at a rich collection of artificial bulk mixtures in terms of variation with regard to cellular composition. This approach is analogous to the pseudo-bulk generation in [11].

In order to compose a single bulk mixture, we took only a single patient from the given single-cell study in order to keep the patient’s cellular *expression* integrity, but we disturbed the cell type-specific *abundance* in order to introduce variance. More precisely, we determined the actual amount of single cells for each cell type within a given patient

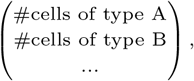

and then perturbed with a normally distributed multiplicative factor given via

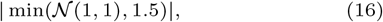

i.e., we arrive at the perturbation

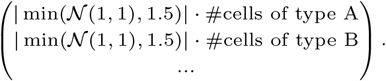

Here we limited to a maximal factor of 1.5 in order to keep oversampling on a moderate scale, but still introduce suitable variance. This approach results in the total amount of cells to be randomly sampled for each cell type out of the given patient. Thus, we arrived at an artificial bulk mixture *Y*_·,*s*_ by summing over all sampled single-cell profiles. Note that here we sampled with replacement, in order to be able to satisfy multiplicative factors larger than one in equation (16). Note that this approach naturally provides us with ground truth cellular proportions, as also explained in [6, Methods] and [11, Supplementary Note 4]. More precisely, the ground truth cell type abundance in bulk sample *m* of cell type *l* can be computed as

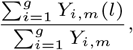

where *Y*_·,*m*_(*l*) denotes the cell type-specific expression of cell type *l* in artifical bulk sample *m* and *Y*_·,*m*_ denotes the total expression of artifical bulk sample *m*.

Representative pseudo-bulk samples and the distribution of cellular abundance across all pseudo bulks are depicted in Figures 11.

##### A.2.4 Gene distortion

In order to exaggerate local inconsistencies, see Section 1, we randomly distorted a fixed amount of genes within the simulated bulk mixtures by a multiplicative factor. This approach is analogous to the linear multiplicative noise model in [11]. Therefore, we fixed the percentage of genes we want to distort in order to arrive at a set 𝒢 of genes to be distorted. In our case we configured this to be 40% of the 1000 genes available. For each gene *g* ∈ 𝒢 we determined a multiplicative distortion factor *ζ*_*g*_ by drawing from the normal distribution,

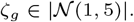

The distribution parameters are chosen to yield biologically reasonable fold changes. This approach provides us with distorted bulk samples by multiplying

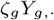

for all *g* ∈ *𝒢*. In particular this means that each gene is distorted constantly across bulk samples in order to simulate technical gene-specific batch effects in gene sequencing measurements.

##### A.2.5 Harp generalizes stably for sufficient amount of training samples

As an initial benchmark we explored Harp’s performance in terms of variable training cohort sizes. Therefore, we generated a total set of 160 bulk mixtures available for training and 40 test bulks. Out of the set of training bulks we subsampled a variable amount of bulk mixtures in the range [6, 160], trained Harp and evaluated its performance on the 40 held-out test samples. The results of this benchmark are depicted in Figure 10. As expected, all quality scores showed consistently inferior results for insufficient training samples and stabilize at competitive values after enough samples were added to the training set. We observed that this stabilization occurred at around 50 training samples, leading to the selection of this cohort size for remaining simulation benchmarks.

**Figure. 10.**
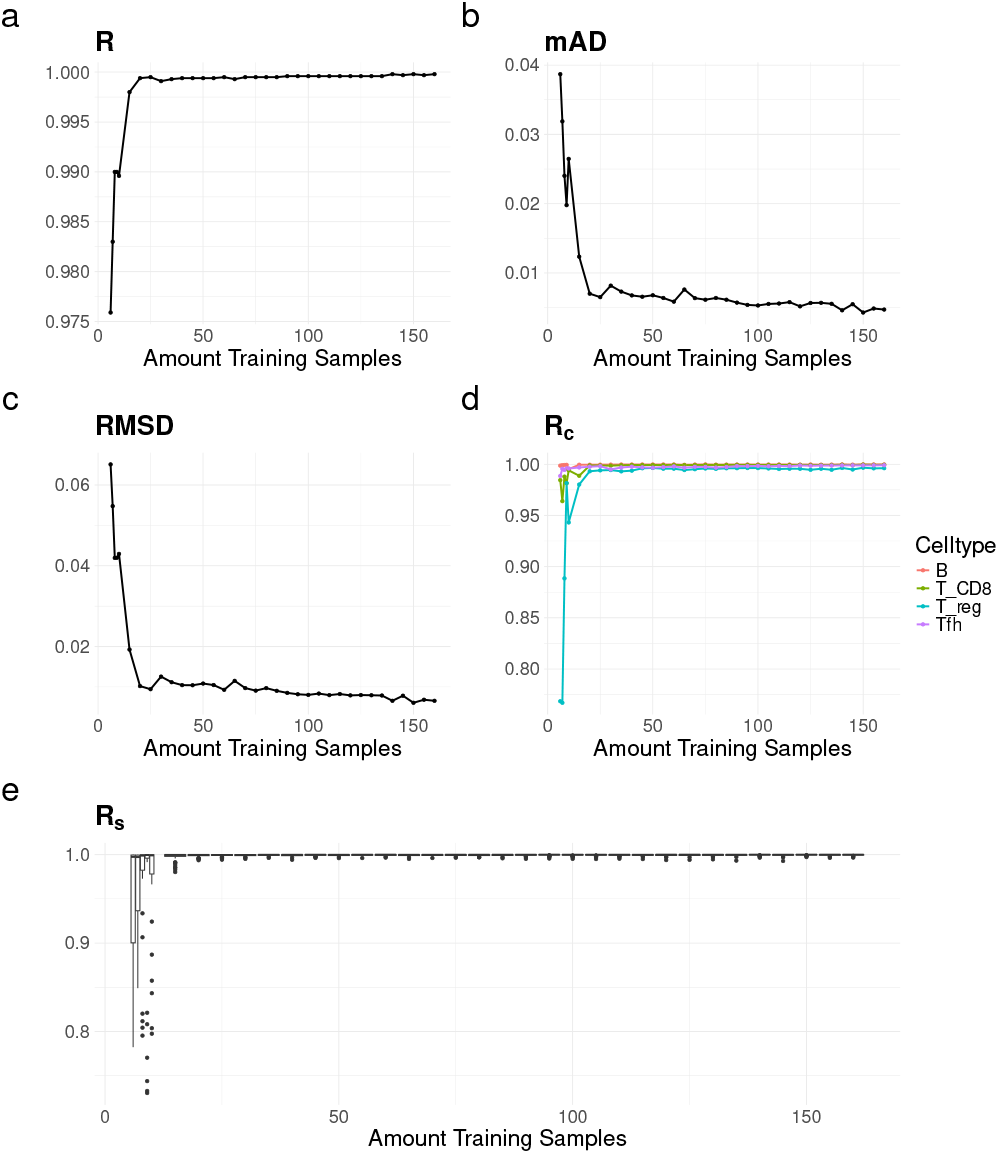
Impact of the amount of artificial bulk samples used for training Harp on main performance metrics.

**Figure. 11.**
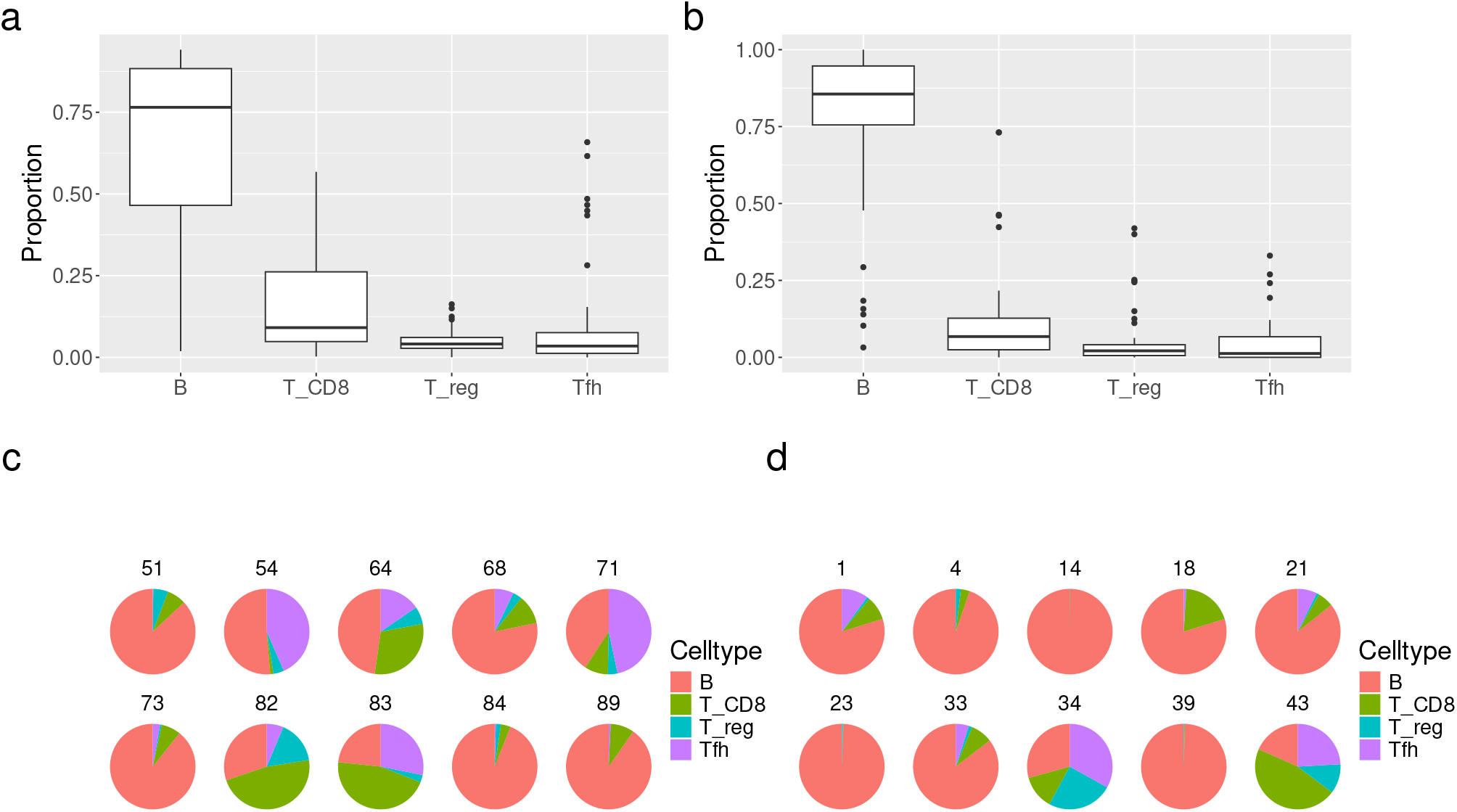
Overview of cellular composition for a selection of test bulk samples (a, c) and training bulk samples (b, d) used in simulations.

**Figure. 12.**
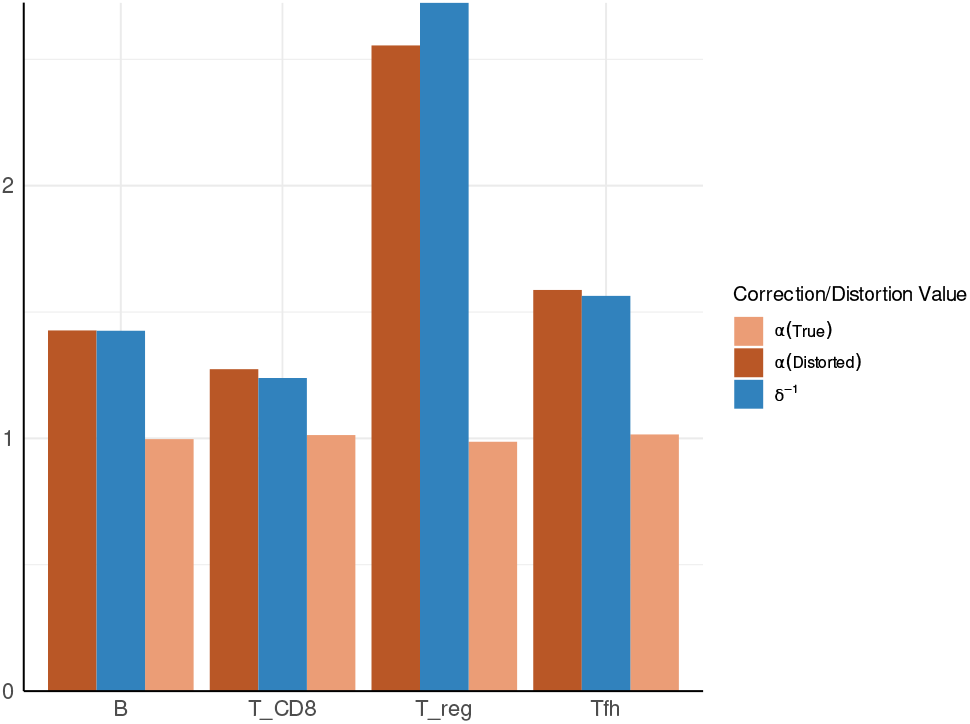
Cell type-specific correction values, ***α***, computed by Harp when provided with correct cell proportions (True) and distorted proportions (Distorted) in *Training* mode, respectively. These values are compared to the inverse distortion rate ***δ***^−1^.

##### A.2.6 Configurations of competing methods for benchmarking deconvolution tools on simulated data

CIBERSORT [27] is designed to use a given reference in deconvolution, and comes with the custom reference LM22 derived from microarray data, see [27]. Its successor, CIBERSORTx [9], provides the option to learn a custom reference from given single-cell or sorted RNA-seq data. Furthermore, CIBERSORTx addresses cross-platform variation via refined batch correction. Therefore, the authors introduced bulk mode (B-mode) and single-cell mode (S-mode) batch correction. In our simulations, as the employed single-cell dataset [24] was generated using 10x Chromium, see Section A.2.1, we followed the authors’ suggestion and applied S-mode batch correction when providing CIBERSORTx with the single-cell dataset. We denote this version of CIBERSORTx as CIBERSORTx (scRNA-seq). For the microarray reference LM22 we followed the authors’ suggestion and applied B-mode batch correction. We denote this version of CIBERSORTx as CIBERSORTx (LM22) in our evaluations. Additionally, instead of the generic microarray reference LM22 we also provided the reference learned by Harp to CIBERSORTx. We note that due to Harp’s capability of adapting its reference to the training bulks we did not apply any of the proposed batch corrections within the CIBERSORTx algorithm on the simulated data in order to underline the fact that our reference already overcame these batch effects by integrating bulk information during training. We denote this approach as CIBERSORTx (Harp).

Note, that when employing LM22 we mapped the granular cell types of this reference to the cell compartments analyzed in simulations as follows: naive and memory B cells were mapped to B cells. Naive, memory resting and memory activated CD4 T cells were mapped to CD 4 T cells.

Concerning BayesPrism, the standard algorithm exploits fine-grained cellular states in order to optimally integrate single-cell information, see [11]. Therefore, we performed automated subclustering interfaced in the Seurat package [33] on the given single-cell library. This provided us with cellular states that refined the original cell types. The subclustering information and the single-cell library were then provided to BayesPrism. This standard usage of BayesPrism is denoted as BayesPrism in our benchmarks. Similarly to CIBERSORTx we also evaluated the performance of BayesPrism when provided with the Harp reference, which is possible as BayesPrism allows for replacing subclustering and single-cell information by directly providing a reference by choosing the input.type = “GEP”, see also the corresponding vignette.^2^ This approach is referred to as BayesPrism (Harp) in the following. However, we note that the recommended default usage of BayesPrism is to receive a rich scRNA-seq count matrix in order to internally derive a reference via maximum likelihood estimation, see [11]. For applying MuSiC [6] we used the default setting as advised in the corresponding documentations. MuSiC does not allow for providing a custom reference as an entire single cell library is required for inference. Note that there exists also a more recent version MuSiC2 [34] which is equivalent to the initial MuSiC implementation in our usecase, as we do not include different sample conditions.

##### A.2.7 Simulating uncertain experimental compositions

As discussed in Section 2.1 flow cytometry measurements suffer from cell type specific bias. Mathematically, we simulated this scenario, by drawing a *cell type-specific distortion rate*, ***δ***(*l*) (with l representing a cell type), from the distribution

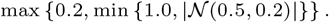

This means, that we limited ***δ***(*l*) in [0.2,1] in order to arrive at biologically reasonable distortions, and allow for a mean loss of 50% within a cell type with a standard deviation of 0.2. These factors were used to simulate compromised proportion measurements via

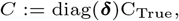

where *C*_True_ is a *q* ×*n* matrix holding the ground-truth cell type proportions for each sample. As expected, Figure 12 reveals, that the cell specific correction rates ***α*** indeed captured the simulated distortion by satisfying the relation ***α***(*l*) ≈ ***δ***^−1^(*l*).

##### A.2.8 Harp’s regularization approach balances deconvolution accuracy and biological integrity

In order to understand Harp’s dependency on the hyperparameter *λ*, see Sections A.1.2 and 2.1, we analyzed its evolution during cross-validation in the *Training* mode of our algorithm and we studied the quality of models fitted for different values of *λ*. Therefore, we used the exact same simulated data as in the main benchmark in Section 3.2.

Concerning the first point, the correlation computed for each candidate *λ* in the cross-validation phase is shown in Figure 13. As expected, we observed optimal performance for an intermediate *λ*^′^ = 0.8. This is due, on the one hand, to the fact that the naive unregularized solution (*λ* = 0) is expected to overfit to the training set and, therefore, cannot optimally explain the held-out samples. On the other hand, excessively high values of *λ* prevent the incorporation of bulk information by sticking to the anchor *X*^*^. This explains why we observed a steep decline after the optimal *λ*^′^ was reached. However, we stress that during cross-validation Harp exploits only a simple least square regression to arrive at a rough estimate of cellular abundance, see Section A.1.2.

**Figure. 13.**
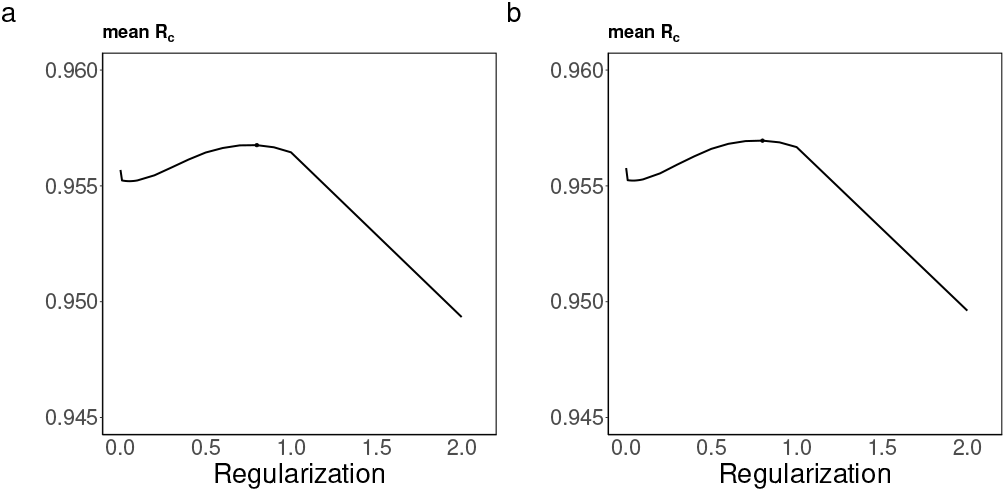
Evolution of *λ* during cross-validation in Harp’s *Training* mode. The optimum is marked as a point. (a) shows the trajectory before cell type-specific correction with *α* and (b) in the second iteration of our proposed updating scheme, i.e., after this correction step.

In Figure 13 (a) we show the trajectory in the first iteration of Harp’s *Training* mode, i.e., where *X*^′^ in equation (4) is determined for ***α*** being the identity matrix. In Figure 13 (b), however, we show the trajectory in the second iteration, i.e., subsequent to determining ***α*** in equation (5). As expected both figures appear identical, because the simulated data uses ground truth proportions for *C* and thus, the ***α*** determined in equation (5) is still an approximate of the diagnonal matrix, see also ***α***(True) in Figure 12.

Next, we had a closer look at the performance of the *entire* Harp algorithm for different *λ* values, see Figure 14. A major difference to the previous paragraph is thus, that DTD is used during *Deconvolution* mode, which is far more elaborate than plain least squares regression. Here, we represent the *R*_*c*_ metric and the *R*_*s*_ metric for varying values of *λ*, see Section 2.2. Regarding cell type-specific performance (*R*_*c*_), we observed that the cell types were generally well captured for suitably small values of *λ* and the optimal lambda value is in the interval [0.5, 0.75] which is a regime consistent with the optimal cross-validated *λ*^′^ = 0.8 from the previous paragraph. However, for values close to zero, *R*_*c*_ behaved instable for regulatory T cells indicating that this regularization regime gives suboptimal deconvolution results. This again underlines that lowly abundant cell compartments are especially challenging in deconvolution, as already noted in Section 3.2. The *R*_*c*_ metric behaved analogously; interestingly, this metric was not prone to instabilities for *λ* values close to zero. This also indicates that tuning *λ* during cross-validation only to optimize the *R*_*c*_ metric, see Section A.1.2, is natural, because the *R*_*s*_ metric is stable independently of *λ*, at least in our simulation scenario. Considering a broader spectrum of *λ* values, Figures 14 (b) and 14 (d) show that both metrics broke down in the regime of very high regularization, indicating that sticking with the naive single-cell reference hinders the adequate integration of bulk information in the *Training* mode of Harp.

**Figure. 14.**
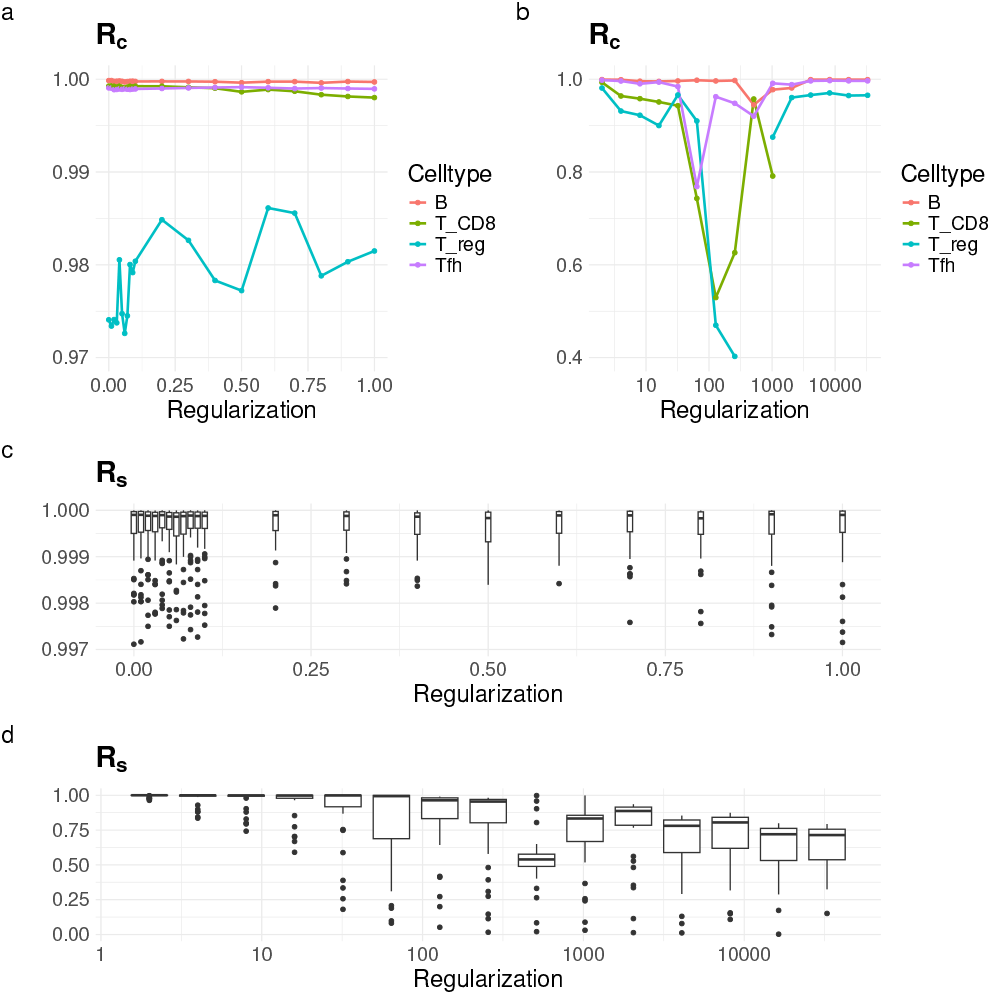
Cell type-specific (a, b) and sample specific performance (c, d) of Harp for varying regularization parameters *λ*, separated in low (a, c), and large regularization (b, d) regimes.

#### A.3 Data harmonization with Harp improved deconvolution accuracy in a study combining data from two distinct sources

##### A.3.1 Data processing

For this case study the bulk gene expression and flow cytometry data were given by [18]. We utilized TPM-normalized gene expression data and cell abundances obtained from BostonGene^3^, provided as part of the Kassandra project [35]. The cell types included in the flow cytometry data were labeled as B Naive, B Ex, B NSM, B SM, Plasmablasts, CD4 T cells, CD8 T cells, Basophils LD, Dendritic cells, Plasmacytoid dendritic cells, Monocytes C, Monocytes I, Monocytes NC, NK cells, B Memory, T cells, Monocytes C+I, Monocytes NC+I, Monocytes, B cells, and Lymphocytes. Since not all cell types were present in every sample, we included only those samples that contained all specified cell types and were paired with bulk RNA-seq data. This filtering resulted in a final dataset comprising 250 samples, which was then divided into a 150-sample training set and a 100-sample test set. The cell types included in the data represented different levels of cell type granularity. For further analysis, we selected B cells (B memory cells and B naive cells), Monocytes, T cells, NK cells, Plasmablasts, Basophils LD, Plasmacytoid dendritic cells, and Dendritic cells. The sum of the cell type populations selected for each sample did not add up to 100%, The cell proportions across samples averaged 95.99%, with values ranging from a minimum of 85.98% to a maximum of 99.98%. Therefore, we added an additional row to the cellular composition matrix to account for the Unidentified cells, see Section A.1.3. To align the flow cytometry data with our references (both sorted RNA-seq and microarray) and/or because of the rarity of certain cell types, we also categorized Basophils LD, Plasmacytoid dendritic cells, and Dendritic cells under the “Unidentified” category in our analysis.

The sorted RNA-seq signature PBMC data was obtained from four healthy donors, as reported in [19], to build the initial reference. To ensure consistency, we mapped or relabeled some of the cell types to align with the reference and flow cytometry dataset. We maintained Plasmablasts as a separate cell type (rather than including them within B cells, despite their small population) because they formed a distinct cluster compared to other B cell subtypes in the reference data, as shown in Figure 15. We then constructed the cell reference matrix *X*^*^ by averaging the profiles of each cell type. For this reference, we retained 1343 intersecting genes shared between the sorted RNA-seq and bulk RNA-seq expression data.

**Figure. 15.**
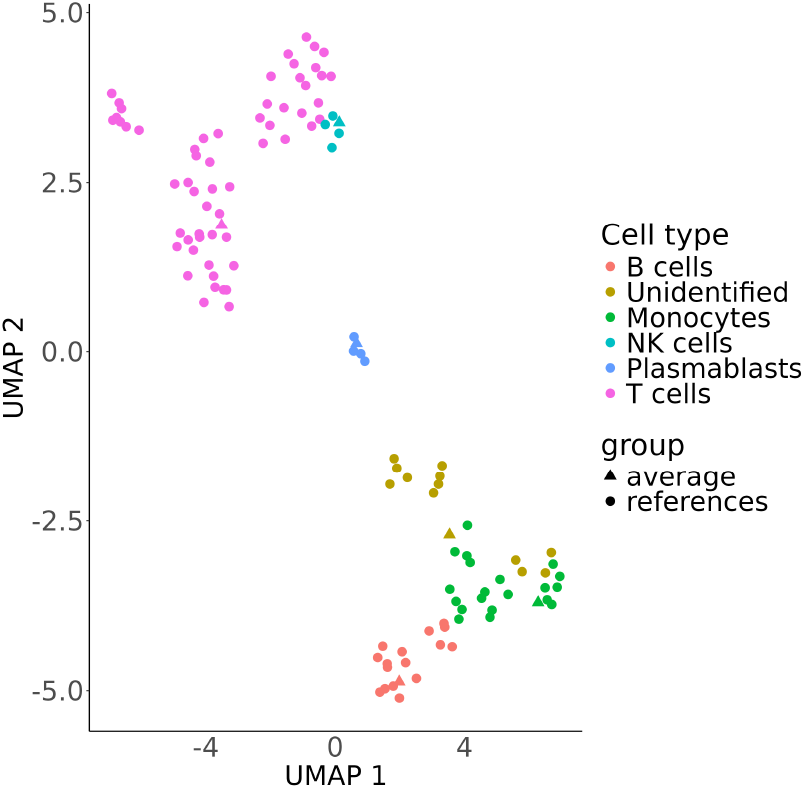
Umap of sorted RNA-seq signature data (after mapping to the cell types corresponding to the flow cytometry data). The triangle marks the average profile of each cell type and each color represents a specific cell type.

Additionally, we incorporated another cell reference dataset derived from microarray technology into our analysis. We utilized LM22, the signature matrix from CIBERSORT [27], which represents 22 distinct PBMC cell types characterized by 547 genes. From these, we selected 503 genes that were shared between the bulk RNA-seq expression data and LM22. Similarly to the other reference, we relabeled certain cells and then computed the average for each cell type.

##### A.3.2 Configurations of competing methods for benchmarking deconvolution tools on RNA-seq data

For benchmarking purposes, we provided BayesPrism and CIBERSORTx with the signature data containing original cell types in both data sets, i.e., sorted RNA-seq and microarray data. After the deconvolution step, we aggregated the proportions of the corresponding cell types, according to the mapping we applied in Section A.3.1, to ensure consistency with the flow cytometry data. This enabled us to calculate the quality scores involving cell abundances. For example, we combined the predicted proportions of B cell subtypes and labeled the aggregated result as “B cells”.

In Figure 6, there is no correlation score for Plasmablasts from CIBERSORT(LM22), because the reference does not include this cell type. Regarding the use of BayesPrism and CIBERSORT(LM22) in our RNA-seq data benchmark, we mainly followed the procedures outlined in Section A.2.6. When providing LM22 to BayesPrism we set the parameter input.type = “GEP”, as only one gene expression signature for each cell type was available. However, it is important to emphasize that BayesPrism is specifically designed to work with scRNA-seq expression data, whereas microarray data is not explicitly supported. For CIBERSORTx (RNA-seq), we enabled B-mode batch correction to deconvolute the bulk samples, using the sorted RNA-seq data as the source GEP (gene expression profile). Batch correction is highly recommended by the authors when the reference signature and bulk samples are measured on different platforms. Therefore, it was not strictly required in this case, as both data sets were measured through RNA sequencing. However, since enabling batch correction improved results for CIBERSORTx (RNA-seq), we decided to retain it. For both imputing cell type fractions and building a signature matrix, quantile normalization was disabled, as the input datasets were not derived from microarray technology. For CIBERSORTx (Harp), we enabled batch correction to be consistent with CIBERSORTx (RNA-seq), but did not select a GEP source in this case. Here, we emphasize that for quality performance regarding cell proportions, we excluded the Unidentified cells from the analysis. We did not include MuSic [6] in these benchmarks because it requires the entire single-cell library as input for the reference, which was not compatible with our reference data. To assess the quality of the reconstructed bulk expression profiles of the algorithms, we calculated the score for the predicted bulk samples for Harp as described in Section 2.2. For CIBERSORT, this score is provided directly in the method’s output, and for BayesPrism, we reconstructed bulk samples by summing the cell expression profiles for each sample, as provided in its output, and then calculated the correlations, *ρ* according to Section 2.2.

##### A.3.3 Evaluation of reconstructed bulk expression profiles of RNA-seq data

With the aim of studying whether the source of cellular composition estimates/measurements or the adjustment of cell references have a greater influence on the accuracy of the reconstructed bulk gene expression profiles, we reconstructed two additional sets of bulk expression data. These reconstructions were similar to the previous ones (discussed in section 4) but used cell abundances derived from Harp this time. In Table 1 and Table 2 the first and third row, with cell proportions labeled “Flow cytometry” were previously discussed in Figure 6 (g) and Figure 7 (g), respectively. Table 1 and 2 show that, in general, as expected, the adjustment of the reference profile (*X* in *Y* = *XC*) has a stronger influence on the quality of bulk expression predictions compared to the source of cell proportions. Comparing two rows with the same reference first and second or third and fourth, implies that the difference in the bulk predictions using different cellular composition matrices (*C* in *Y* = *XC*) is minimal, with only a very small advantage for Harp over the flow cytometry data. Moreover, a comparison of the two tables indicates that using a reference generated with a technology similar to the one the bulk expression data was generated with, in our case sorted RNA-seq reference, provides more accurate results.

**Table 1.**
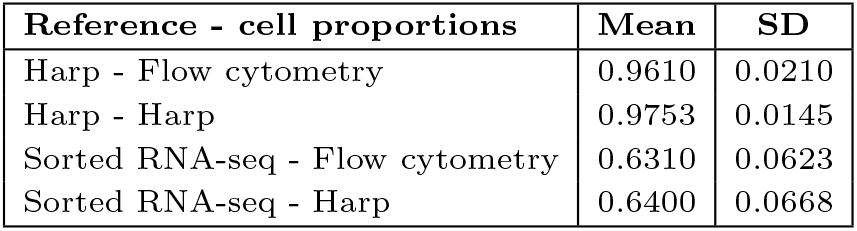
Comparison of reconstructed and observed bulk gene expression in 100 PBMC RNA-seq test samples. The mean and standard deviation of Pearson correlations (*ρ*) were calculated between reconstructed bulk expression profiles and observed RNA-seq data. The reconstructed bulk expression profiles were generated using the Harp reference and sorted RNA-seq data, with cell proportions from Harp and flow cytometry.

**Table 2.** Comparison of reconstructed and observed bulk gene expression in 100 PBMC RNA-seq test samples. The mean and standard deviation of Pearson correlations (*ρ*) were calculated between reconstructed bulk expression profiles and observed RNA-seq data. The reconstructed bulk expression profiles were generated using the reference from Harp and microarray data (LM22), with cell proportions from Harp and flow cytometry.

| Reference - cell proportions | Mean | SD |
| --- | --- | --- |
| Harp - Flow cytometry | 0.8906 | 0.0854 |
| Harp - Harp | 0.9063 | 0.0767 |
| Lm22 - Flow cytometry | 0.5353 | 0.0990 |
| Lm22 - Harp | 0.5660 | 0.1067 |

#### A.4 Harp’s performance remained intact when using data from single microarry source

##### A.4.1 Benchmarking Harp against other deconvolution tools

As a second example using real data, we used 20 PBMC samples from [27], for which bulk gene expression was measured by microarray technology and cell proportions were measured via flow cytometry. For this dataset, we split the samples into a training set consisting of 12 samples and a validation set of eight samples. For the anchor *X*^*^ in Harp we utilized LM22 provided in [27] (for more details see section A.4.4).

Here, we observed that, in general, Harp showed a strong performance, similar to its result discussed earlier. According to the first plot, *R*, in Figure 16a, both Harp and BayesPrism show similar performance, and achieved higher overall correlation compared to CIBERSORT. However, the rest of the plots in Figure 16a show that BayesPrism achieved better RMSD and mAD in comparison to Harp. This could potentially be due to the limited size of Harp’s training set, which is below the threshold derived in simulations, see Section A.2.5. Regarding cell type-specific prediction across samples (*R*_*c*_), as shown in Figure 16c, Harp outperformed both CIBERSORT (LM22) and BayesPrism in inferring the proportions of Memory B cell, CD8 T cells and Monocytes. In contrast, CIBERSORT (LM22) performed the best for Naive B cells and CD4 T cells, while BayesPrism showed an strong performance in predicting NK cell proportions. Figure 16b represents the performance metric *R*_*s*_ and shows that Harp outperformed its competitors in within-sample cell proportion predictions. Harp also achieved better performance than CIBESORT (LM22) in predicting bulk gene expression profiles, see Figure 16d.

**Figure. 16.**
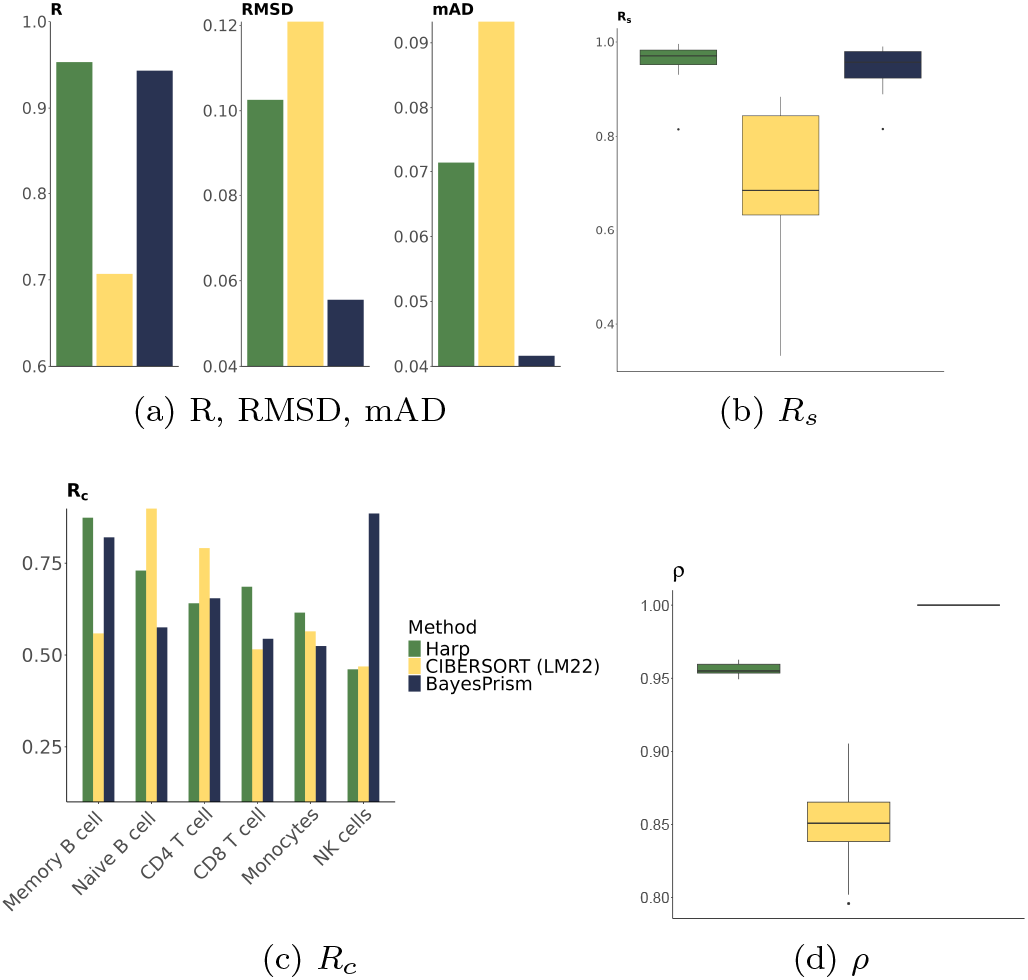
Benchmark of various deconvolution tools (Harp, CIBERSORT and BayesPrism) on eight PBMC microarray bulk expression validation samples with a microarray reference (LM22) using different quality scores. Plots (a-c) evaluate performance on the prediction of cell proportions, while plot (d) analyses the quality of bulk gene expression predictions.

##### A.4.2 Hybrid deconvolution

Similarly to Section 4 and 3.2, we studied the effect of the harmonaized Harp reference (using LM22 as anchor *X*^*^ as in the previous section) on the performance of other tools in deconvolution of the eight PBMC bulk expression validation samples. In general, Figures 17a show that CIBERSORTx benefits from the Harp reference, in contrast to BayesPrism. The Harp reference influences both methods positively in predicting sample specific cell proportions, see Figure 17b. Figure 17c represents that the performance of BayesPrism in predicting proportions of Naive B cells, and CD8 T cells significantly improved when using the Harp reference. However, the predictions for Memory B cells, CD4 T cells and NK cells were negatively affected when using the Harp reference. On the other hand, CIBERSORTx showed better performance with the LM22 rather than the Harp reference, except for CD8 T cells and Monocytes. In terms of bulk gene expression predictions, CIBERSORTx (Harp) outperformed CIBERSORT (LM22) significantly, see Figure 17d. This indicates that Harp successfully compensates for inconsistencies between the cell reference and bulk expression data.

**Figure. 17.**
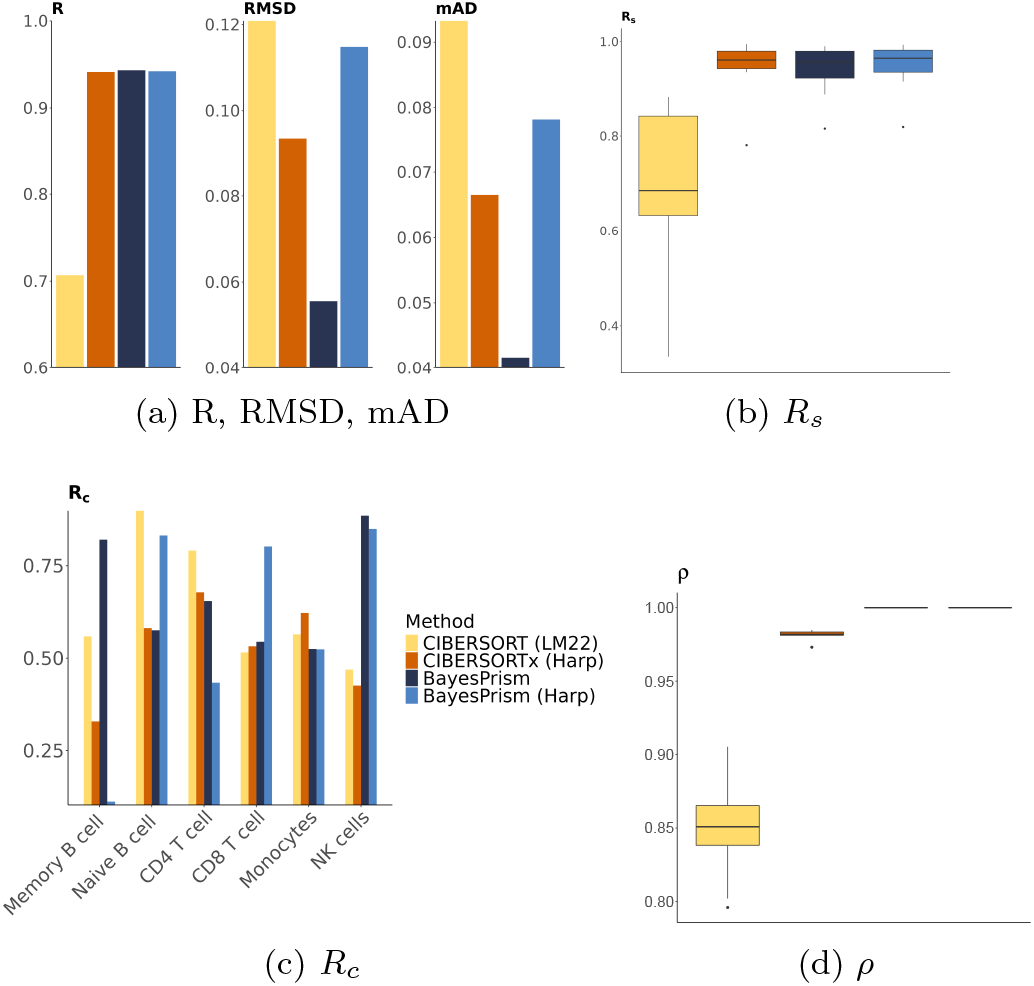
Analysis of the impact of the Harp reference on CIBERSORTx and BayesPrism performance in deconvolution of eight PBMC microarray validation samples, comparing the results with the respective methods using the LM22 reference. Plots (a-d) evaluate performance on the prediction of cell type proportions, while plot (e) analyses the quality of the bulk gene expression predictions.

##### A.4.3 Evaluation of reconstructed bulk expression profiles of microarray data

Analogously to section A.3.3 we studied the effects of different sources of obtained cell proportions and references in the reconstruction of bulk gene expression profiles. Table 3 shows that generally the effect of different references is stronger compared to cell proportions. Moreover, Harp explains the bulk expression data with a correlation of 0.95, showing a major improvement in comparison to the LM22 reference.

**Table 3.** Comparison of reconstructed and observed bulk gene expression in eight PBMC RNA-seq test samples. The mean and standard deviation of Pearson correlations (*ρ*) were calculated between reconstructed bulk expression profiles and observed microarray data. The reconstructed bulk expression profiles were generated using the reference from Harp and microarray data (LM22), with cell proportions from Harp and flow cytometry.

| Reference - cell proportions | Mean | SD |
| --- | --- | --- |
| Harp - Flow cytometry | 0.9397 | 0.0171 |
| Harp - Harp | 0.9560 | 0.0045 |
| Lm22 - Flow cytometry | 0.3598 | 0.0383 |
| Lm22 - Harp | 0.3697 | 0.0291 |

In terms of reconstructing bulk expression data, using the Harp reference and LM22 along cell proportions from flow cytometry, the first and third row in Table 3 (as well as Figure 18, which provides another representation) show that the Harp reference improves the average correlations from 0.35 to 0.93, demonstrating higher accuracy in explaining bulk samples compared to LM22.

**Figure. 18.**
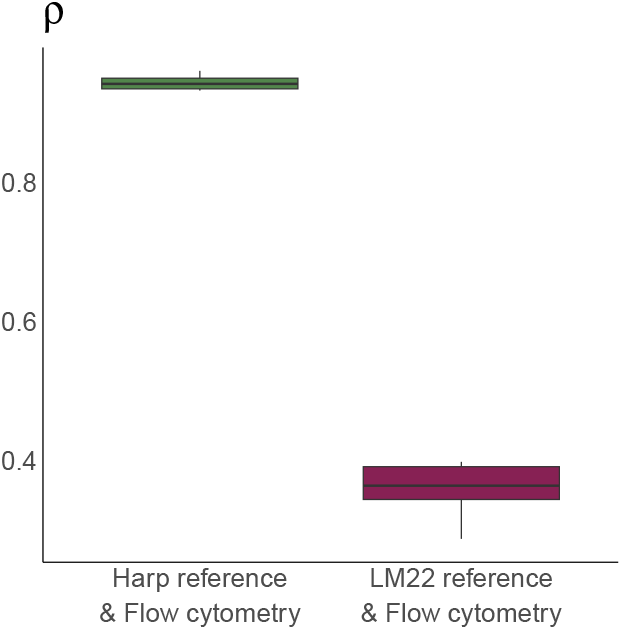
Comparison of the reconstructed bulk gene expression profiles. Box plots show Pearson correlations, *ρ*, between the reconstructed bulk gene expression for eight PBMC microarray samples, using Harp reference (green) and microarray-based reference (LM22) (magenta), and the observed microarray expression data. The cell proportions obtained from flow cytometry data were used for the reconstruction of the bulk samples.

##### A.4.4 Details of data processing of microarray data

The microarray bulk gene expression data included 20 PBMC samples from [27]. For the anchor *X*^*^ in Harp we used the LM22 reference which is the standard CIBERSORT signature [27]. The bulk gene expressions were measured using the Illumina HumanHT-12 V4.0 expression beadchip platform, and the non-normalized data are available on NCBI under the GEO accession number GSE65133. Paired flow cytometry profiling was downloaded from the analyses of [36]. ^4^ The quantified cell populations include Monocytes, Memory B cells, Naive B cells, Gamma Delta T cells, CD4 T cells, CD8 T cells, and NK cells. The proportions of all cell types summed up to 100%, indicating that the data was normalized before. In order to be able to use LM22 as anchor *X*^*^ in our method, we mapped or relabel some of the cell types to align with the flow cytomety data. Here, we did not include the Unidentified row and column in our input data. To utilize bulk expression data in Harp, we first mapped probe IDs to gene names using the R package illuminaHumanv4.db. Next, we calculated the average expression for probes with the same gene symbol within each sample to obtain per-gene expression values. Then quantile normalization was applied to the bulk samples. After splitting the data into training and test set, we trained Harp in its *Training* mode to estimate the Harp reference, we then used it in *Deconvolution* mode to deconvolute bulk samples in the test data set.

##### A.4.5 Configurations of competing methods for benchmarking deconvolution tools on microarray data

To impute cell fractions using CIBERSORT (LM22), we provided CIBERSORT with the gene expression data for all 20 samples, mapped to probe IDs. The data was not aggregated over similar gene names by averaging, allowing CIBERSORT to perform its own method to get the per-gene expression data. Quantile normalization was applied by CIBERSORT, and B-mode batch correction was also enabled, using the LM22 GEP source. Concerning CIBEROSRTx (Harp) and BayesPrism (Harp), we provided CIBERSORTx and BayesPrism with the Harp reference and the bulk samples as was in our test set. BayesPrism was provided with the anchor reference LM22, as described in Section A.3.2. We then calculated performance metrics only for the samples in the test sets.

All algorithms, using different references, predicted negative correlations for Gamma Delta T cells. Therefore, after deconvolution and before calculating the cell proportions related perfomance metrics, we removed this cell type from the analysis. For the reconstructed bulk data evaluation we followed the procedure described in Section A.3.2.

##### Footnotes

1 The preprocessed single-cell data is publicly accessible under 10.5281/zenodo.10139153.

2 The BayesPrism vignette is under https://github.com/Danko-Lab/BayesPrism.

3 The RNA-seq expression and flow cytometry data are also available under the ID sdy67 on BostonGene (https://science.bostongene.com).

4 The flow cytometry data is also available on NCBI, under the GEO accession number of GSE65133, under the ‘Analyze with GEO2R’ section (https://www.ncbi.nlm.nih.gov/geo/geo2r/?acc=GSE65133).

## References

1. Ping Hu, Wenhua Zhang, Hongbo Xin, and Glenn Deng. Single cell isolation and analysis. Frontiers in cell and developmental biology, 4:116, 2016.

2. Regina K. Cheung and Paul J. Utz. Screening: CyTOF-the next generation of cell detection. Nature Reviews Rheumatology, 7(9):502–503, 2011.

3. Angela R Wu, Norma F Neff, Tomer Kalisky, Piero Dalerba, Barbara Treutlein, Michael E Rothenberg, Francis M Mburu, Gary L Mantalas, Sopheak Sim, Michael F Clarke, and Stephen R Quake. Quantitative assessment of single-cell RNA-sequencing methods. Nature methods, 11(1), 2014.

4. H. Kim, N.and Kang, A. Jo, S. Yoo, and H. Lee. Perspectives on single-nucleus rna sequencing in different cell types and tissues. Journal of Pathology and Translational Medicine, 2023.

5. E Denisenko, BB Guo, M Jones, R Hou, L deKock, T Lassmann, D Poppe, O Clément, RK Simmons, R Lister, and ARR. Forrest. Systematic assessment of tissue dissociation and storage biases in single-cell and single-nucleus rna-seq workflows. Genome Biol, 2020.

6. Xuran Wang, Jihwan Park, Katalin Susztak, Nancy R Zhang, and Mingyao Li. Bulk tissue cell type deconvolution with multi-subject single-cell expression reference. Nature Communications, 10(380), 2019.

7. Francisco Avila Cobos, Jo Vandesompele, Pieter Mestdagh, and Katleen De Preter. Computational deconvolution of transcriptomics data from mixed cell populations. Bioinformatics, 34(11):1969–1979, 01 2018.

8. Franziska Görtler, Marian Schoen, Jakob Simeth, Stefan Solbrig, Tilo Wettig, Peter J Oefner, Rainer Spang, and Michael Altenbuchinger. Loss-function learning for digital tissue deconvolution. Journal of Computational Biology, 27(3):342–355, 2020.

9. Aaron M. Newman, Chloé B. Steen, Chih Long Liu, Andrew J. Gentles, Aadel A. Chaudhuri, Florian XScherer, Michael S. Khodadoust, Mohammad S. Esfahani, Bogdan A. Luca, David Steiner, Maximilian Diehn, and Ash A. Alizadeh. Determining cell type abundance and expression from bulk tissues with digital cytometry. Nature Biotechnology, 2019.

10. Franziska Görtler, Malte Mensching-Buhr, Ørjan Skaar, Stefan Schrod, Thomas Sterr, Andreas Schäfer, Tim Beißbarth, Anagha Joshi, Helena U Zacharias, Sushma Nagaraja Grellscheid, and Michael Altenbuchinger. Adaptive digital tissue deconvolution. Bioinformatics, 40(Supplement 1):i100–i109, 06 2024.

11. Tinyi Chu, Zhong Wang, Dana Pe’er, and Charles G. Danko. Cell type and gene expression deconvolution with bayesprism enables bayesian integrative analysis across bulk and single-cell rna sequencing in oncology. Nature Cancer, 2022.

12. Jakob Simeth, Paul Hüttl, Marian Schön, Zahra Nozari, Michael Huttner, Tobias Schmidt, Michael Altenbuchinger, and Rainer Spang. Virtual tissue expression analysis. Bioinformatics, 40(12):btae709, 11 2024.

13. L.X. Garmire, Y. Li, Q. Huang, and et al. Challenges and perspectives in computational deconvolution of genomics data. Nature Methods, 21:391–400, 2024.

14. Grace XY Zheng, Jessica M Terry, Phillip Belgrader, Paul Ryvkin, Zachary W Bent, Ryan Wilson, Solongo B Ziraldo, Tobias D Wheeler, Geoff P McDermott, Junjie Zhu, et al. Massively parallel digital transcriptional profiling of single cells. Nature communications, 8(1):1–12, 2017.

15. Ashraful Haque, Jessica Engel, Sarah A Teichmann, and Tapio Lönnberg. A practical guide to single-cell rnasequencing for biomedical research and clinical applications. Genome medicine, 9(1):1–12, 2017.

16. Jonathan R Chubb, Tatjana Trcek, Shailesh M Shenoy, and Robert H Singer. Transcriptional pulsing of a developmental gene. Current biology, 16(10):1018–1025, 2006.

17. Danson Shek Chun Loi, Lei Yu, and Angela Ruohao Wu. Effective ribosomal rna depletion for single-cell total rna-seq by scdash. PeerJ, 9, 2021.

18. M. T. Zimmermann, A. L. Oberg, D. E. Grill, G. Ovsyannikova, I. H. Haralambieva, R. B. Kennedy, and G. A. Poland. System-wide associations between dna-methylation, gene expression, and humoral immune response to influenza vaccination. PLoS One, 11(3):e0152034, 2016.

19. G. Monaco, B. Lee, W. Xu, S. Mustafah, Y. Y. Hwang, C. Carré, N. Burdin, L. Visan, M. Ceccarelli, M. Poidinger, A. Zippelius, J. Pedro de Magalhães, and A. Larbi. Rna-seq signatures normalized by mrna abundance allow absolute deconvolution of human immune cell types. Cell Reports, 26(6):1627–1640.e7, 2019.

20. E. Brombacher, O. Schilling, and C. Kreutz. Characterizing the omics landscape based on 10,000+ datasets. Scientific Reports, 15:3189, 2025.

21. W Evan Johnson, Cheng Li, and Ariel Rabinovic. Adjusting batch effects in microarray expression data using empirical bayes methods. Biostatistics, 8(1):118–127, 2007.

22. Hung Nguyen, Ha Nguyen, Duc Tran, Sorin Draghici, and Tin Nguyen. Fourteen years of cellular deconvolution: methodology, applications, technical evaluation and Harp outstanding challenges. Nucleic Acids Research, 52(9):4761–4783, 04 2024.

23. Julien Racle, Kaat de Jonge, Petra Baumgaertner, Daniel E Speiser, and David Gfeller. Simultaneous enumeration of cancer and immune cell types from bulk tumor gene expression data. eLife, 6:e26476, 2017.

24. CB Steen, BA Luca, MS Esfahani, A Azizi, BJ Sworder, BY Nabet, DM Kurtz, CL Liu, F Khameneh, RH Advani, Y Natkunam, JH Myklebust, M Diehn, AJ Gentles, AM Newman, and AA. Alizadeh. The landscape of tumor cell states and ecosystems in diffuse large b cell lymphoma. Cancer Cell, 2021.

25. T Roider, J Seufert, A Uvarovskii, F Frauhammer, M Bordas, N Abedpour, M Stolarczyk, JP Mallm, SA Herbst, PM Bruch, H Balke-Want, M Hundemer, K Rippe, B Goeppert, M Seiffert, B Brors, G Mechtersheimer, T Zenz, M Peifer, B Chapuy, M Schlesner, C Müller-Tidow, S Fröhling, W Huber, S Anders, and S. Dietrich. Dissecting intratumour heterogeneity of nodal b-cell lymphomas at the transcriptional, genetic and drug-response levels. Nature Cell Biology, 2020.

26. Leland McInnes, John Healy, and James Melville. Umap: Uniform manifold approximation and projection for dimension reduction. arXiv preprint 1802.03426, 2018.

27. Aaron M. Newman, Chih Long Liu, Michael R. Green, Andrew J. Gentles, Weiguo Feng, Yue Xu, Chuong D. Hoang, Maximilian Diehn, and Ash A. Alizadeh. Robust enumeration of cell subsets from tissue expression profiles. Nature Methods, 12(5):453–457, 2015.

28. Anupama Reddy, Jenny Zhang, Nicholas S. Davis, Andrea B. Moffitt, et al. Genetic and functional drivers of diffuse large B cell lymphoma. Cell, 171(2):481–494.e15, 2017.

29. Roland Schmitz, George W. Wright, Da Wei Huang, Calvin A. Johnson, et al. Genetics and pathogenesis of diffuse large B-cell lymphoma. New England Journal of Medicine, 378(15):1396–1407, 2018.

30. Jorge Nocedal and Stephen J Wright. Numerical optimization. Springer, 1999.

31. Justine Hsu, Julien Jarroux, Anoushka Joglekar, Juan P. Romero, Corey Nemec, Daniel Reyes, Ariel Royall, Yi He, Natan Belchikov, Kirby Leo, Sarah E.B. Taylor, and Hagen U Tilgner. Comparing 10x genomics single-cell 3’ and 5’ assay in short-and long-read sequencing. bioRxiv, 2022.

32. Jakob Simeth, Paul Hüttl, Marian Schön, Zahra Nozari, Michael Huttner, Tobias Schmidt, Michael Altenbuchinger, and Rainer Spang. Virtual tissue expression analysis. Bioinformatics, page btae709, 2024.

33. Tim Stuart, Andrew Butler, Paul Hoffman, Christoph Hafemeister, Efthymia Papalexi, William M Mauck, Yuhan Hao, Marlon Stoeckius, Peter Smibert, and Rahul Satija. Comprehensive integration of single-cell data. cell, 177(7):1888–1902, 2019.

34. Jiaxin Fan, Yafei Lyu, Qihuang Zhang, Xuran Wang, Mingyao Li, and Rui Xiao. Music2: cell-type deconvolution for multi-condition bulk rna-seq data. Briefings in Bioinformatics, 23(6):bbac430, 2022.

35. Aleksandr Zaitsev, Maksim Chelushkin, Daniiar Dyikanov, Ilya Cheremushkin, Boris Shpak, Krystle Nomie, Vladimir Zyrin, Ekaterina Nuzhdina, Yaroslav Lozinsky, Anastasia Zotova, Sandrine Degryse, Nikita Kotlov, Artur Baisangurov, Vladimir Shatsky, Daria Afenteva, Alexander Kuznetsov, Susan Raju Paul, Diane L. Davies, Patrick M. Reeves, Michael Lanuti, Michael F. Goldberg, Cagdas Tazearslan, Madison Chasse, Iris Wang, Mary Abdou, Sharon M. Aslanian, Samuel Andrewes, James J. Hsieh, Akshaya Ramachandran, Yang Lyu, Ilia Galkin, Viktor Svekolkin, Leandro Cerchietti, Mark C. Poznansky, Ravshan Ataullakhanov, Nathan Fowler, and Alexander Bagaev. Precise reconstruction of the tme using bulk rna-seq and a machine learning algorithm trained on artificial transcriptomes. Cancer Cell, 40(8):879–894.e16, August 2022.

36. Francesco Vallania, Andrew Tam, Shane Lofgren, Steven Schaffert, Tej D. Azad, Erika Bongen, Winston Haynes, Meia Alsup, Michael Alonso, Mark Davis, Edgar Engleman, and Purvesh Khatri. Leveraging heterogeneity across multiple datasets increases cell-mixture deconvolution accuracy and reduces biological and technical biases. Nature Communications, 9(1):4735, 2018.

